# Within-plant variation in chemical defence of *Erysimum cheiranthoides* does not explain *Plutella xylostella* feeding preference

**DOI:** 10.1101/2024.11.01.621493

**Authors:** Kunqi Wang, Tobias Züst

## Abstract

Plants invest a substantial fraction of their resources into defence against herbivores, with the highest levels of defence often allocated only to the most valuable tissues. Plants in the genus *Erysimum* (Brassicaceae) evolved the ability to produce novel cardenolides in addition to ancestrally conserved glucosinolates. While plants co-express both defences, differences in tissue-specific expression might represent an effective cost saving strategy.
Larvae of the glucosinolate-resistant diamondback moth *Plutella xylostella* occasionally feed on *Erysimum cheiranthoides* but tend to avoid younger leaves. Here, we predicted that caterpillar feeding preference is shaped by variation in cardenolide levels, thus we quantified within-plant variation in defensive and nutritional traits of vegetative or early reproductive plants and performed feeding assays to evaluate the relative importance of cardenolides.
In accordance with optimal defence theory (ODT), youngest leaves contained the most nutrients and the highest levels of cardenolides, glucosinolates, and trichomes, with more extreme within-plant differences found in reproductive plants. Caterpillars consistently avoided the well-defended youngest leaves both on whole plants and with detached leaf discs. Surprisingly, neither experimental addition (by external application) nor removal (by CRISPR-Cas9 knockout) of cardenolides significantly affected caterpillar feeding preference.
Physical and chemical defences, including cardenolides, co-vary within *E. cheiranthoides* to maximise defence of youngest leaves. While *P. xylostella* clearly responds to some of these traits, the prominent cardenolide defence appears to lack potency against this specialist herbivore. Nonetheless, the careful regulation and re-mobilization of cardenolides to younger leaves during plant development suggests an important role for plant functioning.

**One-sentence summary:** Distribution of ancestral and novel defensive chemicals *Erysimum cheiranthoides* vary within plant according to relative leaf value and leaf age, but in isolation, chemical defences are insufficient to affect feeding preference of *Plutella xylostella* larvae.

## Introduction

Plants protect themselves against enemy attack using multiple defensive traits, including physical structures and chemical compounds (Gong & Zhang 2014). Defence traits are generally considered to be costly for the plant as they divert resources away from growth and reproduction (Züst *et al*. 2011; Züst & Agrawal 2017). While resource-based allocation costs can be context-dependent and not always apparent (Koricheva 2002), it is nonetheless clear that plant investment in defence must be substantial. For example, total investment in defensive glucosinolate production by *Arabidopsis thaliana* was estimated to be at least 15% of all photosynthetic energy (Bekaert *et al*. 2012). In line with these theoretical predictions, well-defended genotypes of *A. thaliana* with high glucosinolate concentrations and trichome densities had lower fruit numbers compared to less-defended genotypes when grown in a field experiment where herbivores had been artificially removed (Mauricio & Rausher 1997; Mauricio 1998). Similarly, knockout mutants of *A. thaliana* with abolished accumulation of aliphatic and indole glucosinolates, or with abolished trichome production showed increased growth relative to wildtype plants, suggesting that these knockout mutants could benefit from defence-related cost savings (Züst *et al*. 2011).

Costs of defence also can explain patterns in allocation of defences within individual plants as described by the Optimal Defence Theory (ODT). If defence is costly and resources are limited, plants are expected to allocate most of their defences to tissues that are most valuable for fitness, e.g., their youngest leaves for vegetative plants, and their flowers and seeds for reproductive plants (McKey 1974). The ODT has been extensively tested and validated, with many studies finding the highest concentration of chemicals found in high-value younger tissues, regardless of the type of defence or experimental conditions (Traw & Feeny 2008; McCall & Fordyce 2010; Keith & Mitchell-Olds 2017). Consistent with the ODT, *A. thaliana* accumulates the highest levels of glucosinolates in its youngest leaves, followed by a relocation of glucosinolates to flowers and seeds after the initiation of flowering (Brown *et al*. 2003).

Despite their substantial investment in defence and its optimised distribution, most plants are nonetheless susceptible to specialist herbivores with evolved tolerance mechanisms against these defences. For example, plants in the Brassicaceae are commonly attacked by a diverse community of insect herbivores (Bidart-Bouzat & Kliebenstein 2008; Mertens *et al*. 2021), many of which have specialized adaptations against the glucosinolate defence (Winde & Wittstock 2011; Jeschke *et al*. 2017). Potentially in response to this specialist threat, plants in the diverse Brassicaceae genus *Erysimum* evolved the ability to produce cardenolides as a ‘second line’ of chemical defence, and co-express them with the ancestrally conserved glucosinolates (Züst *et al*. 2018, 2020). Glucosinolates are secondary metabolites found in all members of the Brassicaceae, functioning by enzyme-mediated release of highly reactive isothiocyanates and related toxic compounds (Halkier & Gershenzon 2006). In contrast, cardenolides sporadically occur across twelve botanical families and function as specific inhibitors of animal Na^+^/K^+^ ATPase, resulting in impaired nerve signal transduction and herbivore paralysis (Agrawal *et al*. 2012). Importantly, specialized herbivore adaptations for coping with cardenolides are unrelated to glucosinolate tolerance (Agrawal *et al*. 2012), suggesting that Brassicaceae herbivores had no pre-adaptations for coping with cardenolides when these compounds first evolved in the *Erysimum* system approximately 2-5 million years ago (Züst *et al*. 2020).

Cardenolides are potent defences against many herbivores, but likely carry substantial metabolic cost as well. In the common milkweed *Asclepias syriaca*, the best-studied cardenolide system, foliar cardenolide concentrations are negatively correlated with early plant growth across a set of natural genotypes (Züst *et al*. 2015). Furthermore, constitutive and induced concentrations generally show negative correlations across multiple milkweed species (Agrawal & Hastings 2019), suggesting the existence of allocation-based limitations to cardenolide expression. Across the *Erysimum* genus, ancestral glucosinolates and novel cardenolides are co-expressed in most species (Züst *et al*. 2020), likely imposing substantial metabolic costs on the plants. Interestingly, while glucosinolates are at least partially inducible by exogenous jasmonic acid application in most *Erysimum* species, cardenolides are only expressed constitutively (Züst *et al*. 2020). Their co-expression and differential regulation by herbivore cues thus strongly suggest that the two chemical defences evolved in response to distinct evolutionary drivers, serve distinct functions, and may show distinct within-plant allocation patterns.

Here we examine the within-plant distribution of the main defensive traits of the annual plant *E. cheiranthoides* and test their effects on *Plutella xylostella* as one of the main remaining *Erysimum* herbivores. When given free choice, larvae of *P. xylostella* preferentially feed on the lowest leaves while avoiding the uppermost, youngest leaves on the plant. We hypothesized that the glucosinolate-resistant *P. xylostella* is deterred by cardenolides, which according to ODT we expected to be highest in the youngest leaves. We first performed a detailed profiling of within-plant differences in concentrations of glucosinolates and cardenolides on four-week and six-week-old plants to capture the peak vegetative and early reproductive growth stages, and further related these to trichome density and leaf nutrient content. We then performed a series of choice and performance experiments, first confirming 1) *P. xylostella* preference by releasing caterpillars on whole plants, followed by quantification of the location of damage. Next, we compared this preference to 2) performance when caterpillars were caged to leaves of four age classes (approximated by vertical position within the plant). To further isolate potential mechanisms in caterpillar preference and performance, we also tested 3) caterpillar choice and performance on *E. cheiranthoides* leaf discs cut from leaves of four age classes to exclude the effects of plant architecture. Finally, we tested 4) caterpillar choice and performance on broccoli leaf discs painted either with crude extracts from *E. cheiranthoides* leaves from four age classes, or with pure defence compounds at corresponding concentrations, to evaluate the effects of leaf metabolites in isolation.

## Materials and Methods

### Plant and insect material

Seeds of *Erysimum cheiranthoides* accession ‘Elbtalaue’ (Arabidopsis Biological Resource Center accession CS29250, National Plant Germplasm System accession PI 691911) were used for all experiments. The seeds for this accession had originally been collected in Germany and had been propagated by self-fertilization for 7-8 generations in a greenhouse. We also used one CRISPR/Cas9 knockout mutant with fully abolished cardenolide production, but intact glucosinolate production in the same genetic background, allowing us to further test the mechanistic role of cardenolides in herbivore feeding preference (Arabidopsis Biological Resource Center accession CS29906, Younkin *et al*. 2024).

For germination of experimental plants, seeds of *E. cheiranthoides* were soaked in water and kept at 4 °C for three days to break dormancy. Then, three soaked seeds were placed on the soil of pots (7 × 7 × 8 cm) filled with a 1:1 mixture of washed sand and peat-based planting substrate (Einheitserde, Patzer Erden GmbH, Germany). Pots were covered with transparent plastic hoods and placed in a growth chamber (16:8 h light: dark, 23 °C, 60% RH). After one week, hoods and extra seedlings were removed to leave only one seedling per pot. Plants were grown in the growth chamber for a total of three weeks and then moved to a greenhouse (natural light supplemented artificially to 16 h light, 23 °C). Plants were grown in the greenhouse for an additional 1 or 3 weeks, resulting in four-week or six-week-old plants, respectively. Six-week-old plants had initiated flower bud production, but no plants were flowering at the time of experiments. For all assays requiring leaves of age classes, we counted all leaves on a plant and then marked leaves at the 20th, 40th, 60th and 80th percentile along the tip-to-base axis of each plant, labelling them as age classes L4 (lowest/oldest) to L1 (highest/youngest).

An inbred laboratory strain of *Plutella xylostella* was used for all herbivore experiments (see supplementary methods for rearing details). To set up batches of caterpillars for experiments, we either used fresh (laid within the last 24h) or refrigerated eggs to set up additional diet cups and performed experiments with caterpillars in the L2 or L3 stage 8-10 days later (Supplementary Table S1).

### 1. Plant traits

#### 1.1 Sample collection and plant secondary metabolite quantification

To quantify variation in chemical profiles within plants, we sampled leaves of four age classes on six four-week and six-week-old plants each. To account for potential additional variation within leaves, we sampled each leaf at 2-6 positions by cutting out small leaf discs (5 mm diameter; 2 plant ages x 6 replicates x 4 leaf ages x 6 sampling positions = 213 discs). Each disc was added to a 2 mL screw-cap vial containing 50 µL 70% MeOH with three internal standards (ouabain, digitoxin, and sinigrin, 2 ug/mL each), and four 3 mm steel beads. Leaf discs were extracted on a FastPrep shaker (MP Biomedicals, France) at 6.5 m/s for 10 sec, followed by centrifugation at 2×10^4^ g for 5 min and 4 °C. We transferred 40 µL supernatant to a new 1.5 mL tube and centrifuged this sample a second time at 2×10^4^ g for 5 min and 4°C to ensure the absence of any particles. Finally, 30 µL of the twice-centrifuged supernatant were transferred to a glass vial fitted with a glass micro insert for analysis by UPLC-MS.

The relative abundance and richness of cardenolides were analysed by UPLC-MS, using a Vanquish^TM^ Horizon UHPLC System (Thermo Fisher Scientific, USA) coupled to a timsTOF Pro mass spectrometer (Bruker Daltonics, Germany). Plant extracts were separated on an Acquity Premier CSH C18 2.1 x 100 mm column with 1.7 µm pore size (Waters, USA), maintained at 40°C. For detection of cardenolides, 1 µL of the extract was injected and separated at a constant flow rate of 0.55 mL/min with a gradient as follows: 0-4.5 min from 15% to 65% B, 4.5-4.7 from 65% to 100% B, followed by a 1.5 min wash phase at 100% B and 1.7 min reconditioning at 15% B. The mass spectrometer was operated in positive ionization mode at 4,000 V capillary voltage and 500 V endplate offset with an N2 nebulizer pressure of 2.8 bar and dry gas flow of 8 min at 220 °C. Spectra were acquired in the mass range from m/z 50 to 2,000 at ca. 20,000 resolution (m/z 622) and a 1.5 Hz rate. The mass analyser was calibrated at the beginning of each LC run using a 10 mM solution of sodium formate that was injected using a 6-port-valve with a 20 μL loop, giving a mass accuracy below two ppm. We putatively identified cardenolides by their exact mass, fragments, and retention times, and we then integrated peak areas of characteristic mass signals for relative quantification of compounds across all samples (Supplementary Table S2).

For absolute quantification of cardenolides, a subset of five plant samples spanning the full range of cardenolide concentrations were re-analysed by HPLC-UV-MS on an Agilent 1260 Infinity II LC system (Agilent, Switzerland) with UV detection, coupled to an Agilent InfinityLab LC/MSD single quadrupole mass spectrometer with electrospray ionization. Samples were separated on an InfinityLab Poroshell EC-C18 3 x 150 mm column with 2.7 µm pore size (Agilent, Switzerland). The column was maintained at 40 °C and injections of 5 µL were eluted at a constant flow rate of 0.5 mL/min with a gradient of 0.1% formic acid in water (A) and 0.1% formic acid in acetonitrile (B) as follows: 0-16 min from 20% to 28% B, 16-21 min from 28% to 65% B, 21-22.5 min from 65% to 100% B, followed by a 5 min wash phase at 100% B, and 7.5 min reconditioning at 20% B. Cardenolides were quantified using the UV signal at 220 nm, and identified by matching UV peaks to corresponding mass signals monitored using single ion recording. UV peak areas of quantified cardenolides were converted to absolute concentrations using a digitoxin concentration series (20, 100, 200, 300 µg/mL), assuming equivalent UV absorption by equimolar amounts of cardenolides (Züst *et al*. 2019). We then used the concentrations of cardenolides determined by UV signals in the re-analysed samples to convert MS peak areas into absolute concentrations for all samples and compounds.

Glucosinolates were first identified and quantified by UPLC-MS using an equivalent method as for cardenolides, with the mass spectrometer operating in negative ionization mode. However, due to higher concentrations and stronger ionization of glucosinolate compounds, we experienced substantial issues with detector saturation. We therefore only relied on quantification by UV and re-analysed a complete set of samples (2 samples per plant/leaf age = 96 samples) by HPLC-UV-MS. Samples were separated on an InfinityLab Poroshell CS-C18 3 x 100 mm column with 2.7 µm pore size (Agilent, Switzerland). The column was maintained at 40 °C and injections of 3 µL were eluted at a constant flow rate of 0.5 mL/min with a gradient of 0.1% formic acid in water (A) and 0.1% formic acid in acetonitrile (B) as follows: 0-7 min from 2% to 30% B, 7-10 min from 30% to 45% B, 10-10.5 min from 45% to 100% B, followed by a 4.5 min wash phase at 100% B, and 5 min reconditioning at 2% B. Glucosinolates were quantified using the UV signal at 229 nm, and identified by matching UV peaks to corresponding mass signals monitored using single ion recording (Supplementary Table S2). UV peak areas of quantified glucosinolates were adjusted by published response factors to account for differences in compound-specific absorption (Grosser & van Dam 2017) and then converted to concentrations using a sinigrin concentration series (5, 25, 50, 100, 200 µg/mL).

#### 1.2 Trichome quantification

We quantified trichomes on the leaves of three six-week-old *E. cheiranthoides* plants. Leaves of four age classes were marked on each plant as above. For each marked leaf, we acquired high-resolution images of the abaxial and adaxial side using an Emspira 3 digital microscope (Leica Microsystems, Switzerland). Trichomes were then counted on a defined area using ImageJ (Schneider *et al*. 2012) to quantify trichome densities.

#### 1.3 Macronutrient determination

Nutrient content was quantified from a separate set of 18 six-week-old plants. Leaves of four age classes were marked as above, and the marked leaf and one adjacent leaf were harvested, flash-frozen in liquid nitrogen, and later freeze-dried for 48h. Leaves from three plants were pooled to ensure sufficient material for all assays, leaving six biological replicates. Soluble sugar and starch concentration of leaves was determined enzymatically (Velterop & Vos 2001; Smith & Zeeman 2006; Machado *et al*. 2013). Dried leaves were broken up and 10 mg of leaf material per biological replicate was added to a 2 mL screw-cap tube with three steel beads and ground on a FastPrep shaker at 6.5 m/s for 10 sec. Next, we added 500 µL 80% (v/v) EtOH (Fisher Chemical) to each tube, followed by incubation on a thermomixer at 78 °C for 15 min and 800 rpm, and centrifugation at 2×10^4^ g for 10 min at 4 °C. Supernatants were transferred to a new tube, and the precipitated pellet was re-extracted with 500 µL 50% (v/v) EtOH (Fisher Chemical). Supernatants were combined and used for sugar determination, while the remaining pellet was used for starch determination.

For starch digestion, we added 1 mL of ddH2O to each sample tube to suspend the remaining pellet. We pipetted 100 µL of this suspension to individual wells of a 2 mL deep well plate. As a control for the starch digestion reaction, we included six wells of pure starch (Sigma-Aldrich, Switzerland) at different concentrations on each deep plate (100 µL of 0.1, 0.5, 1, 1.6, 2.4, and 3 mg of starch in 500 µL ddH2O). We then added 500 µL of a fructose solution (55 mg fructose in 250 mL water) to all wells containing sample or starch suspensions to serve as an internal standard. We closed the plates tightly with a sealing mat and wrapped them in aluminium foil before autoclaving them for 1h. After the plates had cooled to room temperature, we added 500 µL of a digestive enzyme solution to each well, consisting of 50 mL of a 50 mM sodium acetate buffer at pH 5.5 (Sigma-Aldrich, Switzerland), 1.35 mL amyloglucosidase (1.9 U/reaction, Roche, Switzerland) and 12 µL α-amylase (3 U/reaction, Sigma-Aldrich, Switzerland). Plates were then shaken and incubated overnight at 37 °C.

Glucose, fructose, sucrose, and glucose from digested starch were determined by a spectrophotometer using the absorption of NADPH that is produced from the enzymatic conversion of glucose-6-phosphate to 6-phosphogluconate. We prepared a master mix consisting of 20 mL 50 mM HEPES (Sigma-Aldrich, Switzerland) with 5 mM MgCl_2_ (Sigma-Aldrich, Switzerland) at pH 7, 12 mg NADP (Sigma-Aldrich, Switzerland), 20 mg ATP (Sigma-Aldrich, Switzerland) and 20 µL glucose-6-phosphate dehydrogenase (0.14 U/reaction, Roche, Switzerland). We combined 200 µL master mix and 25 µL of sample in wells of a 96-well microplate and recorded NADPH baseline absorbance at 340 nm. We then sequentially added a series of enzyme solutions to quantify the different sugars. First, we added 5 µL hexokinase (40 µL in 600 µL HEPES buffer, 0.5 U/reaction, Roche, Switzerland) to convert glucose to glucose-6-phosphate (and fructose to fructose-6-phosphate). Second, we added 5 µL phosphoglucoisomerase (40 µL in 600 µL HEPES buffer, 1.2 U/reaction, Roche, Switzerland) to convert fructose-6-phosphate to glucose-6-phosphate. Finally, we added 5 µL of invertase (48 mg in 600 µL HEPES buffer, 120 U/reaction, Sigma-Aldrich, Switzerland) to convert sucrose into glucose and fructose. After each addition of an enzyme, we monitored absorbance at 340 nm until a stable state was reached. We quantified all sugars relative to a glucose concentration curve (0, 0.125, 0.25, 0.5, 1 and 2 mM) which was included on each microplate.

Total protein content of the same leaves was quantified using a protocol by Clissold et al. (2006) with slight modifications. In brief, total proteins were extracted from 10 mg freeze-dried plant powder using 0.5 mL 0.1M NaOH with sonication for 30 min and heating in 90°C water bath for 15 min. After centrifugation and transfer of supernatants to a new tube, extraction was repeated with 0.3 mL 0.1M NaOH. The two supernatants were pooled and neutralized using 13 µL 6M HCl, after which we precipitated proteins using 90 µL trichloroacetic acid (100% w/v, Sigma-Aldrich, Switzerland). After removal and discarding of supernatants, the remaining protein pellets were resuspended in 1 mL 0.1M NaOH and diluted 1/20 with water. We determined protein concentrations from three technical replicates using QuickStart^TM^ Bradford Dye (BioRad, Switzerland) by quantifying absorbance at 595 nm on a photospectrometer. Absolute concentrations were calculated using a standard curve of immunoglobulin G (Sigma-Aldrich, Switzerland; 0, 1, 2, 4, 6, 8, 12 µg protein per assay).

### 2. Whole plant choice and performance assay

Whole plant choice assays with *P. xylostella* were performed in the same greenhouse as was used for growing plants. Nine four-week and six-week-old plants each were individually set up in cages made from fine mesh (24.5 x 24.5 x 63 cm, Bugdorm, MegaView Science Co., Taiwan). We then released 20 larvae of *P. xylostella* (10-day-old, Supplementary Table S1) at the base of each plant. After 24h of feeding, every second leaf from the base to the tip of each plant was removed, mounted on sheets of paper, and scanned at 300 dpi on a CanoScan LiDE 220 scanner for quantification of the total leaf area and leaf area consumption using the software ImageJ (Schneider *et al*. 2012). We later repeated this assay with a CRISPR/Cas9 cardenolide knockout mutant, using three six-week-old mutant and wildtype plants each.

No-choice performance assays were performed in an incubator (16:8 h light: dark, 23°C, and 60% RH). Using nine four-week and six-week-old plants each, we marked leaves of four age classes. We then weighed individual caterpillars (10-day-old, Supplementary Table S1) to the nearest 0.01 mg on a microbalance (Ohaus Explorer™ Semi-Micro, Ohaus Waagen, Switzerland) and caged individual caterpillars to each marked leaf using a clip cage (2 plant ages x 9 replicates x 4 leaf ages = 72 cages). Clip cages consisted of two foam rings (5 cm outer diameter, 3.5 cm inner diameter) with a fine mesh glued to one side. Foam rings were placed on the upper and lower sides of a leaf and fixed together with metal clamps. After 24h of feeding, caterpillars were removed and weighed again, and the damaged leaves were scanned to determine leaf area consumption.

### 3. Leaf disc choice and performance assay

Leaves of different ages differ in shape and size, with at least some caterpillars having clear preference of feeding locations within-leaf (Shroff *et al*. 2008). Furthermore, attached leaves may respond to herbivore feeding by induction of plant defences (even though defence inducibility of *E. cheiranthoides* was found to be low, Züst *et al*. 2020). To exclude differences in leaf size and reduce the potential of induced changes, we performed a choice assay in an incubator (16:8 h light: dark, 23 °C, 60% RH) using detached leaf discs. Based on our results of the whole plant choice assay, differences among leaves of different ages were more distinct for older plants, thus we only used six-week-old plants for this assay. We marked leaves of four age classes on a new set of 10 plants. From each plant, we cut 2 sets of leaf discs using a cork borer (7.6 mm diameter), using the marked leaves and one adjacent leaf each. We set up a total of 20 Petri dishes (5 cm diameter) as choice arenas by adding a thin layer of 3% agar to each dish and placing one disk per leaf age at each of the corners of a 2 x 2 cm square on top of the agar. We then added one *P. xylostella* larva (10-day-old, Supplementary Table S1) in the centre of each petri dish. The Petri dishes were sealed with Parafilm to prevent the escape of caterpillars. After 24h, all caterpillars were removed, and discs were scanned to determine leaf area consumption.

We also performed a no-choice performance assay using leaf discs from six-week-old *E. cheiranthoides* plants. Leaves of four age classes were marked on a set of six plants as above and used to cut a total of 12 leaf discs (14.9 mm diameter) per leaf age. For the youngest leaves (L1), some leaves were too small to cut a leaf disc, thus we used entire leaves instead. We filled wells of 12-well plates with a thin layer of 1% agar and added leaf discs of each leaf age to the wells in a randomized pattern. We then added one pre-weighed larva of *P. xylostella* (10-day-old, Supplementary Table S1) to each well (12 replicates per leaf age). Plates were sealed with Parafilm to prevent the escape of caterpillars. After 24h, caterpillars were weighed again, and the remaining leaf discs were scanned to determine leaf area consumption.

### 4. Leaf extract and pure compound assays

Leaves differ in their physical properties, nutritional content, and secondary metabolite concentrations. To further narrow down the traits responsible for the differential preference of *P. xylostella* larvae, we performed choice and performance assays in which we applied crude *E. cheiranthoides* leaf extracts to broccoli leaf discs (see supplementary methods). Broccoli (*Brassica oleracea*) is a preferred food plant for *P. xylostella* that is low in glucosinolates and free of trichomes. We also applied commercially available glucosinolate or cardenolide compounds to broccoli leaf discs to evaluate their effect on *P. xylostella* feeding preference in isolation. For glucosinolates, we used sinigrin (Sigma-Aldrich, Switzerland) or a mixture of glucoiberin and glucocheirolin (Phytolab GmbH, Germany), while for cardenolides we used digitoxin or convallatoxin (both Sigma-Aldrich, Switzerland). Sinigrin and digitoxin are both commonly used reference compounds, glucoiberin and glucocheirolin are the main glucosinolates of *E. cheiranthoides*, and convallatoxin is an isomer of identified compound stro.2 (Supplementary Table S2) and representative of other *E. cheiranthoides* compounds in terms of polarity. We dissolved each compound or compound mixture in 100% MeOH and diluted them to concentrations quantified in L4 to L1 leaves of six-week-old plants. We then applied these four concentrations to sets of broccoli leaf discs (7.6 mm diameter) in Petri dish choice arenas and added one larva of *P. xylostella* (9-day-old, Supplementary Table S1) to the centre of each dish. After 24h, all caterpillars were removed, and discs were scanned to determine leaf area consumption.

### Statistical analysis

All analyses were performed in the statistical software R v4.0.4 (R Core Team 2021). Concentrations or relative abundances of cardenolide or glucosinolate compounds were analysed with separate linear mixed effects models (function lme in the nlme package; Pinheiro *et al*. 2021). For each compound, we fit the log-transformed abundances quantified from individual leaf discs to fixed effects of leaf age, plant age, and their interaction. Leaf identity nested within plant identity was specified as random effects to account for repeated sampling of multiple discs from the same leaves (technical repeats), and of multiple leaves from the same plants (leaf ages). We extracted means and 95% confidence intervals from these models using function predictSE from the AICcmodavg package (Mazerolle 2020). Differences in concentrations between four-week and six-week-old plants for each leaf age were assessed using pairwise linear contrasts (function emmeans; Lenth 2022).

Leaf trichomes of six-week-old plants were analysed using a mixed effects model, fitting trichome density to fixed effects of leaf age, leaf side (adaxial/abaxial), and their interaction. Leaf identity nested within plant identity was specified as random effect to account for repeated measurements of the same plant (technical repeats and leaf ages). Leaf nutrient concentrations of six-week-old plants were analysed by equivalent mixed effects models, using fixed effects of leaf age and random effects of plant identity to account for repeated measurements of the same plant (leaf ages). For trichome and nutrient models, we analysed untransformed data, but optimized model fit by estimating separate variances for each leaf age (function *varIdent* in the nlme package).

To analyse plant damage caused by free-moving caterpillars on whole plants, we first fit separate LOESS smoothers of leaf area damage against continuous leaf numbers, numbered bottom to top for each plant. We then used these smoothers to extract the predicted damage at the 20^th^, 40^th^, 60^th^ and 80^th^ leaf percentile (corresponding to leaf ages L4 to L1) for each plant and analysed these using a linear mixed effects model. Log-transformed damage was fit to fixed effects of leaf age, plant age, and their interaction, and plant identity was specified as random effect. To analyse performance of caterpillars constrained to specific leaves using clip cages, we calculated relative growth rates of each caterpillar using

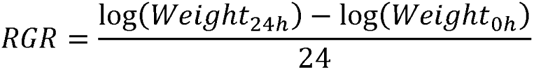

We then analysed caterpillar growth rate and the log-transformed consumed leaf area by separate linear models using generalized least squares (function *gls* in package nlme), fitting leaf age and plant age as explanatory variables, and allowing for separate variances for each leaf age.

To analyse caterpillar choice in leaf disc assays, we fit a mixed effects model of consumed leaf area to fixed effects of leaf age with a random effect of replicate identity, to account for non-independence of leaf discs from the same choice assay replicate. To analyse caterpillar performance in no-choice leaf disc assays, we fit relative growth rate and consumed leaf area as a function of leaf age using linear models as above.

## Results

### 1. Plant traits

We putatively identified and quantified 16 cardenolide compounds and two glucosinolates in the leaves of *E. cheiranthoides* (Supplementary Table S2). Both groups of defensive compounds varied substantially with leaf age and plant age. Averaged across all leaves (and leaf ages), four-week-old plants contained 1.66 ± 0.36 µmol cardenolides g^-1^ dry weight (mean ± 1 SE) and 7.88 ± 1.38 µmol glucosinolates g^-1^ dry weight. In six-week-old plants, these concentrations increased by approximately 70% each, to 2.80 ± 0.36 µmol cardenolides g^-1^ dry weight and 13.40 ± 1.38 µmol glucosinolates g^-1^ dry weight. Differences between four-week and six-week-old plants were even more pronounced when leaf age was considered (Figure 1).

**Figure 1.**
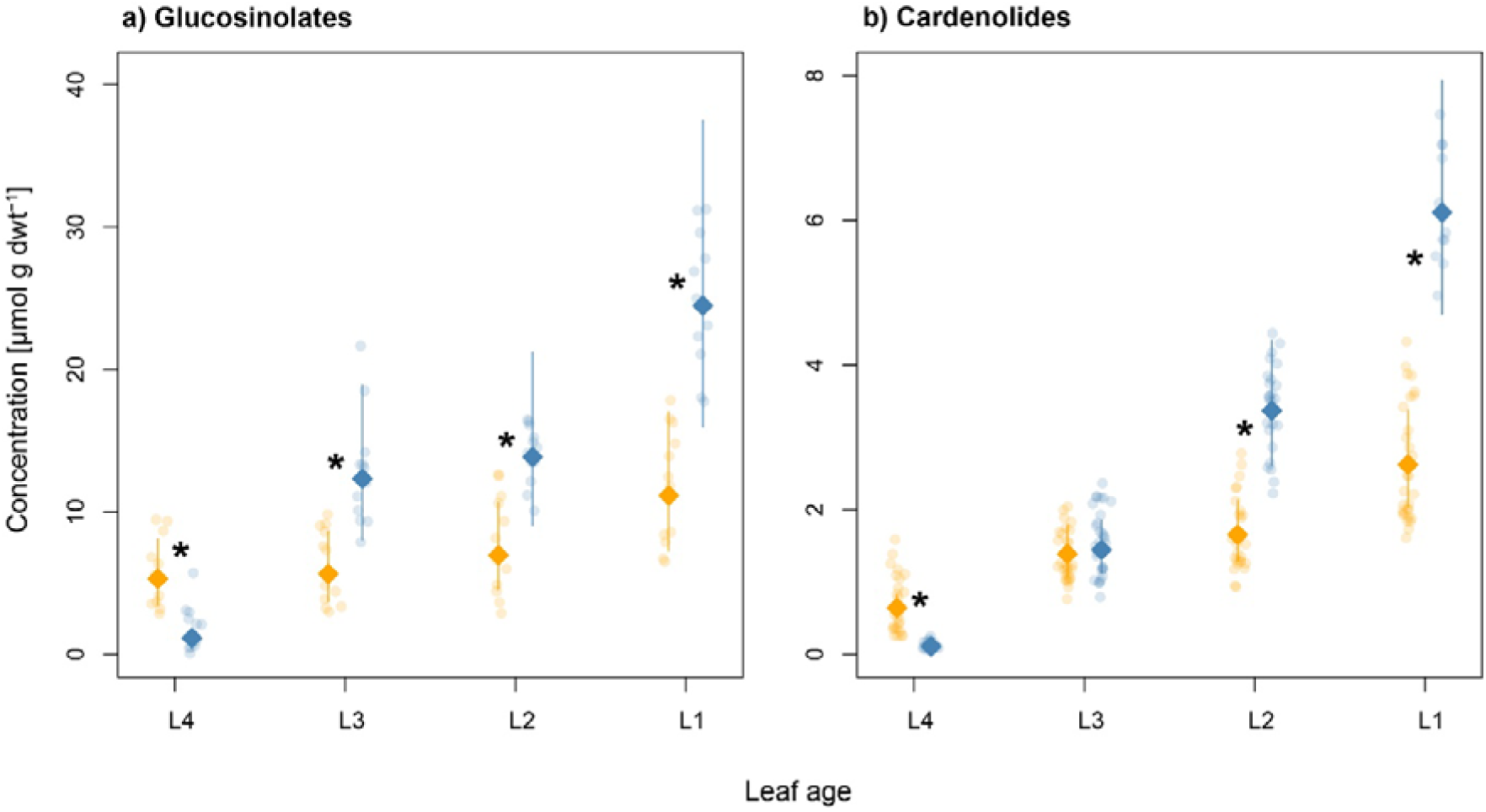
Absolute concentrations [µmol g dwt^-1^] of a) total glucosinolate and b) total cardenolide compounds in leaves of *E. cheiranthoides* from four age classes of four-week (orange symbols) and six-week-old plants (blue symbols). Points are values from individually quantified leaf discs, while diamonds and lines are the estimated mean concentrations and 95% confidence intervals from separate mixed effects models. Asterisks (*) indicate significant pairwise age differences between leaves of the same leaf age.

For four-week-old plants, total cardenolide concentrations differed 4-fold between the oldest and youngest measured leaves (L4: 0.64 µmol g^-1^, L1: 2.63 µmol g^-1^), while glucosinolate concentrations only differed 2-fold (L4: 5.31 µmol g^-1^, L1: 11.14 µmol g^-1^). In six-week-old-plants, within-plant differences increased to 55-fold for cardenolides (L4: 0.11 µmol g^-1^, L1: 6.11 µmol g^-1^) and 20-fold for glucosinolates (L4: 1.14 µmol g^-1^, L1: 24.48 µmol g^-1^), both due to substantially increased concentrations in the youngest leaves and decreasing concentrations in the oldest leaves (Figure 1). Interestingly, individual compounds varied distinctly with plant age (Supplementary Figure S1). For example, the most abundant compound erychroside constituted up to 50% of total cardenolides in the youngest leaves on four-week-old plants, but its absolute concentrations did not significantly increase with plant age. Instead, the increased concentrations in cardenolides of six-week-old plants was primarily due to increases in erysimoside, erycordin, and various lower-concentrated compounds (Supplementary Figure S1). The youngest leaves on six-week-old plants thus not only had higher total cardenolide concentrations, but significantly shifted cardenolide profiles as well.

In addition to chemical defences, we also quantified trichome density and nutritional quality of six-week-old plants, given their more dramatic within-plant differences compared to four-week-old plants. As with chemical defences, we found trichome densities to be different among leaves of different ages (F_3,6_ = 199.15, p < 0.001), and to also vary slightly between leaf sides (leaf age × leaf side: F_3,32_ = 2.962, p = 0.047). The youngest leaves on six-week-old plants had more than 3-fold higher trichome densities compared to the oldest leaves (Figure 2a). All leaf nutrients differed significantly with leaf age. Soluble monosaccharides glucose (F_3,15_ = 35.57, p < 0.001) and fructose (F_3,15_ = 11.06, p < 0.001) had the highest concentrations in the oldest leaves and decreased in concentration towards the tip of the plant (Figure 2b-c). Sucrose (F_3,15_ = 190.68, p < 0.001) and digestible starch (F_3,15_ = 88.25, p < 0.001) had relatively homogeneous distributions across leaves of different ages, except for substantially reduced concentrations in the oldest leaves (Figure 2d-e). Finally, total hydrolysable proteins (F_3,15_ = 36.09, p < 0.001) exhibited the same gradient in concentration as leaf defensive traits, with the youngest leaves having 1.8-fold higher concentration compared to the oldest leaves (Figure 2f).

**Figure 2.**
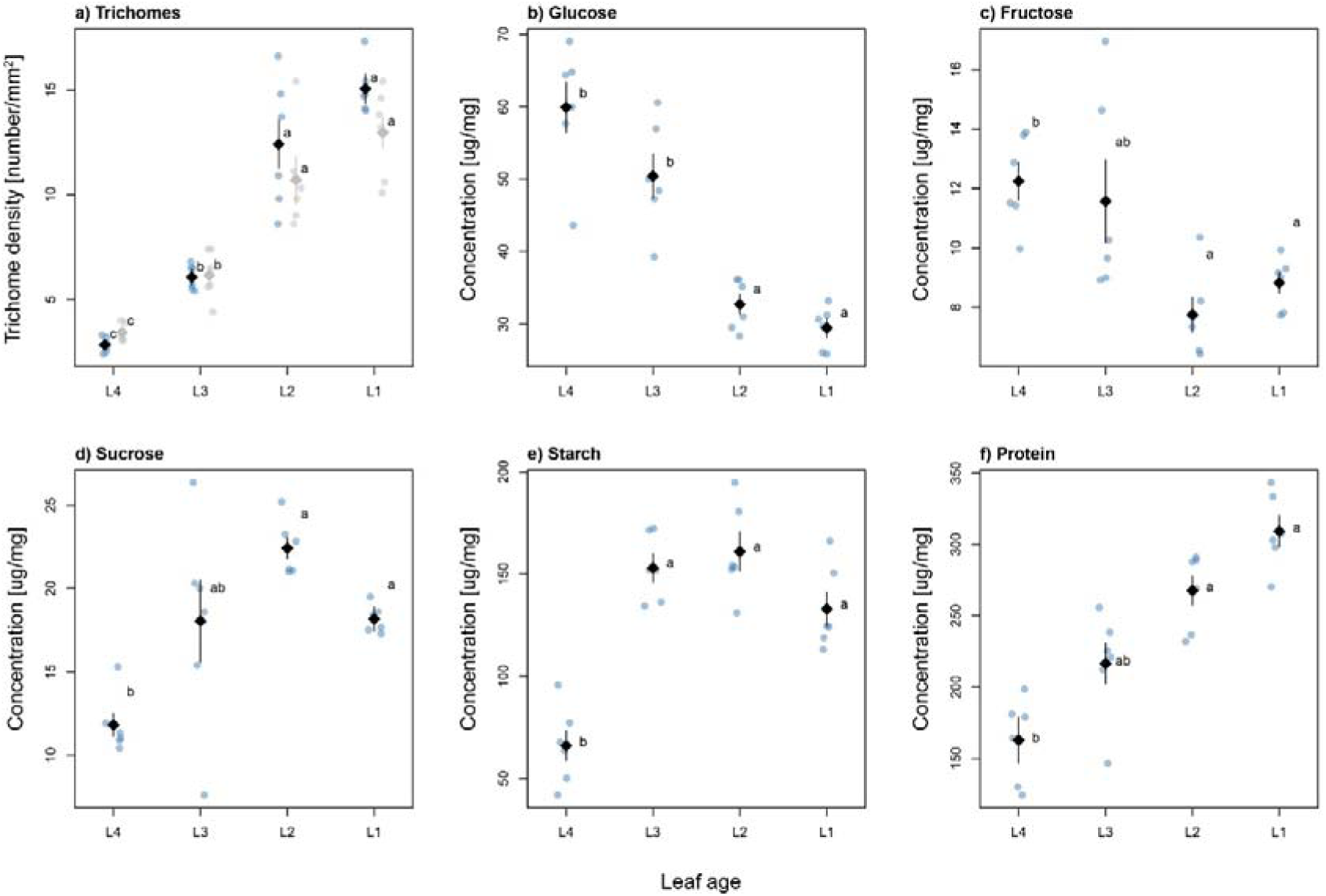
Nutrient content and trichome densities of leaves from four age classes of six-week-old *E. cheiranthoides* plants. a) Trichome densities on the adaxial (blue/black symbols, left) and abaxial (grey symbols, right) leaf side. b-d) Concentration of soluble sugars, d) digestible starch, and e) total hydrolysable protein content. Points are individual measurements, while diamonds and lines are means and 95% confidence intervals from a linear model. Letters denote significant linear contrasts at the p<0.05 level.

### 2. Whole plant choice and performance assay

When larvae of *P. xylostella* were given the choice of where to feed on whole plants, both four-week and six-week old plants were predominantly damaged on their lower, older leaves, with younger leaves receiving the least damage (Figure 3a-b). Four-week-old plants had fewer leaves, and damage was distributed relatively evenly, whereas for the many more leaves of six-week-old plants, damage was concentrated on the lower third. When estimating mean damage at four leaf age classes, we found that damage on four-week-old plants between the oldest and youngest measured leaves only varied 2-fold, whereas it varied over 6-fold on six-week-old plants (leaf age × plant age: F_3,48_ = 5.57, p = 0.002). We repeated this assay using CRISPR/Cas9 knockout plants, but surprisingly, we found no difference in herbivore feeding preference between cardenolide-free mutants and cardenolide-producing wildtype plants (Supplementary Figure S2, leaf age × genotype: F_3,12_ = 0.12, p = 0.947).

**Figure 3.**
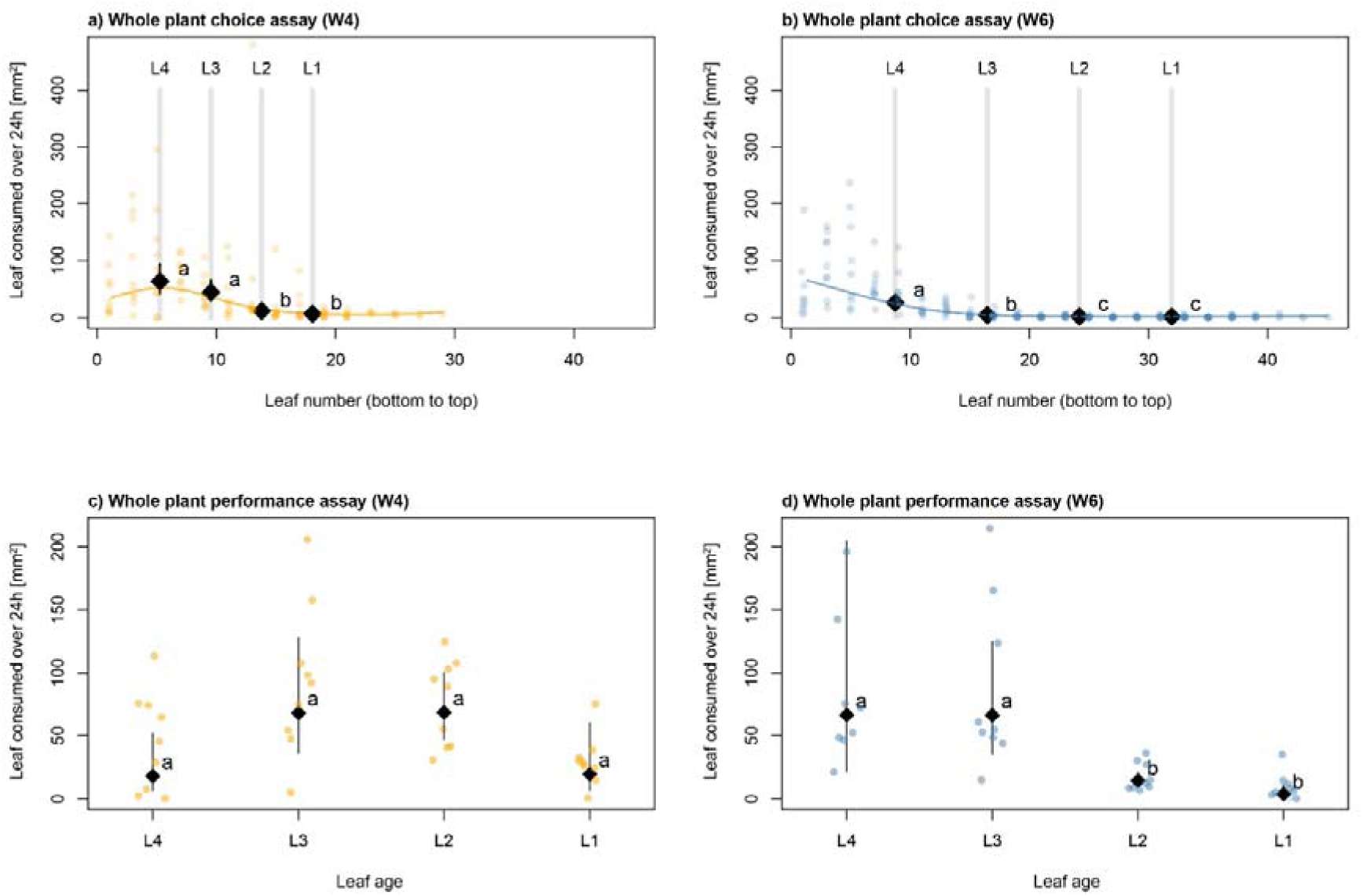
Leaf area consumption by *P. xylostella* under choice and no-choice conditions on whole plants. a-b) Damaged leaf area (mm^2^) on four-week (W4) and six-week-old (W6) plants (n=9 each) after 24 hours of feeding by 20 free-moving caterpillars. Points are individual leaves, while coloured lines are LOESS smoothers to visualize damage patterns. Grey vertical lines and numbers indicate the position of leaf age classes used in other assays, based on leaf number percentiles (20^th^, 40^th^, 60^th^ and 80^th^ percentile). Diamonds and lines are the mean damage and 95% confidence intervals, estimated for these leaf ages from mixed effects models. Letters denote significant linear contrasts at the p<0. 05 level. c-d) Damaged area (mm^2^) on four-week (W4) or six-week-old (W6) plants (n=9 each) after 24 hours of feeding by individual caterpillars confined to one leaf using clip cages. Diamonds and lines are the estimated mean damage areas and 95% confidence intervals from a linear model.

We next quantified caterpillar performance on whole plants by attaching individual caterpillars to leaves of a specific age on four-week or six-week-old plants using clip-cages. After 24 hours of feeding, we quantified weight gain and leaf area consumed by these caterpillars. Caterpillar weight gain, quantified as relative growth rate (RGR), differed significantly by leaf age and plant age (F_3,63_ = 3.45, p = 0.022), with caterpillars restricted to younger leaves having reduced growth or even losing weight on four-week and six-week-old plants (Supplementary Figure S3). Total leaf consumption mirrored these results (Figure 3c-d), and again significantly differed by leaf age and plant age (F_3,63_ = 6.22, p < 0.001). For both measures of caterpillar performance, differences among leaf ages were more pronounced on older plants. On four-week-old plants, caterpillars only grew marginally worse when caged on the youngest leaves (Supplementary Figure S3a) and did not differ in leaf area consumption (Figure 3c). In contrast, caterpillars grew significantly worse on the youngest leaves of six-week-old plants (Supplementary Figure S3b) and consumed up to 18-fold less leaf material on the youngest leaves relative to the oldest (Figure 3d).

### 3. Leaf disc choice and performance assay

To exclude potential effects of plant architecture and defence induction, we next performed choice and performance assays using cut leaf discs. Given the weak effects we found on four-week-old plants, we only used six-week-old plants for these assays. Caterpillars that were given the choice between leaf discs from four age classes consumed significantly more area on older leaves (Figure 4a, F_3,57_ = 23.90, p < 0.001). Similarly, caterpillars that were constrained to feed on one leaf age class consumed significantly more leaf material on older leaves (Figure 4b, F_3,57_ = 15.83, p < 0.001). However, while growth rate of these caterpillars was likewise affected by leaf age (F_3,43_ = 5.59, p = 0.003), only caterpillars feeding on the second-oldest leaves (L3, Figure S4) had a marginally increased growth rate, while caterpillars feeding on the oldest leaves did apparently not benefit from their increased consumption.

**Figure 4.**
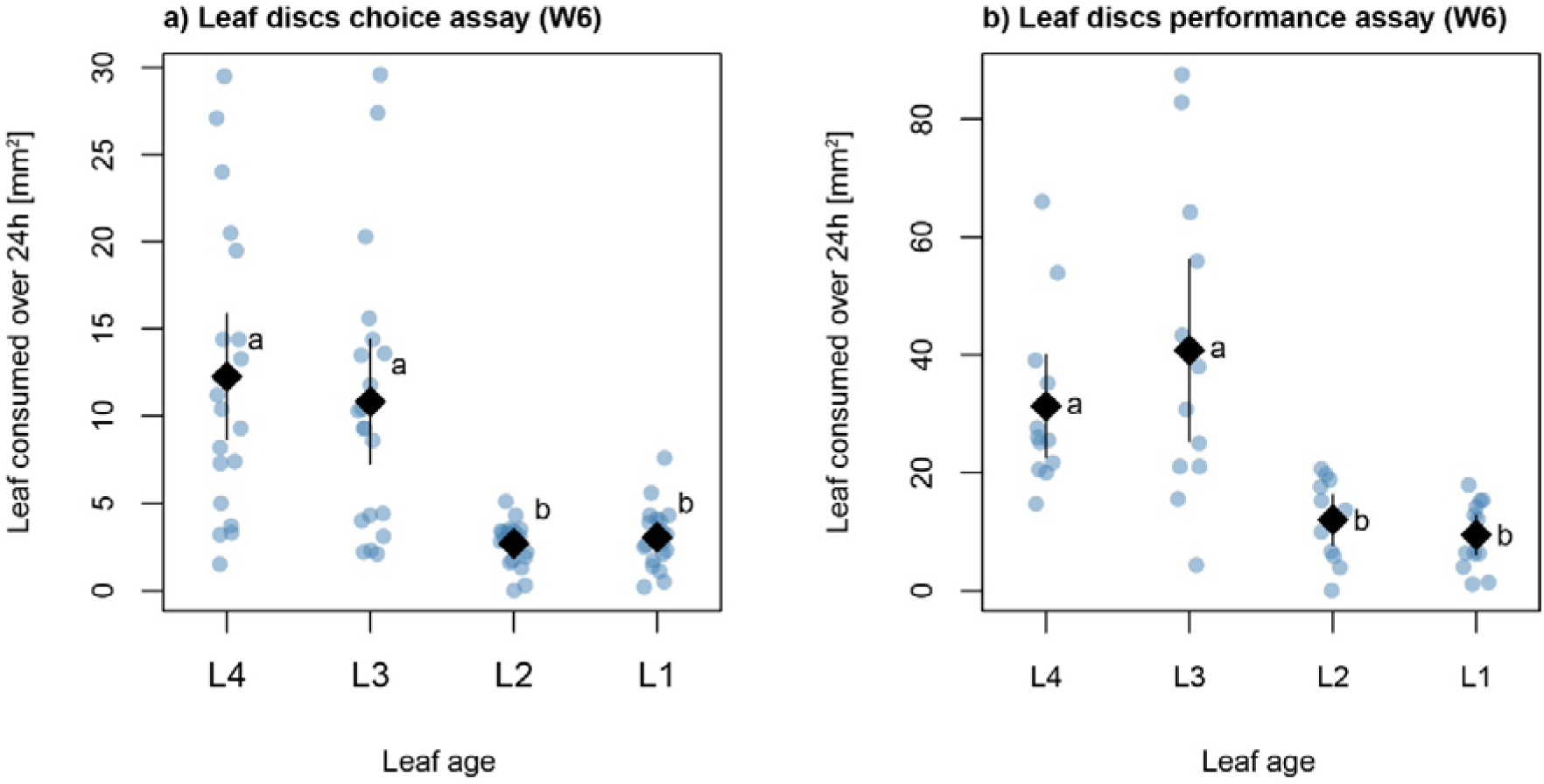
Leaf area consumption by *P. xylostella* under choice and no-choice conditions on detached leaf discs from six-week-old plants (W6). a) Damaged areas on leaf discs of a four-way choice assay (n=20) after 24 hours of feeding by a single caterpillar. Points are values from individual leaf discs, while diamonds and lines are the estimated mean damage areas and 95% confidence intervals from a mixed effects model. Letters denote significant linear contrasts at the p<0.05 level. b) Damaged areas on leaf discs of a no-choice assay (n=12 per leaf age) after 24 hours of feeding by a single caterpillar. Diamonds and lines are the estimated mean damage areas and 95% confidence intervals from linear models.

### 4. Leaf extract and pure compound assays

To exclude effects of nutritional quality and physical defences, we performed choice and performance assays using *E. cheiranthoides* crude leaf extracts painted onto broccoli leaf discs. Caterpillars were given a choice between a control disc painted with solvent and four discs painted with leaf extracts from four age classes. Leaf damage was significantly affected by the extract treatment (F_4,88_ = 16.52, p < 0.001), indicating a slight preference for solvent controls over leaf extracts (Supplementary Figure S5). In no-choice performance assays, neither leaf consumption (Supplementary Figure S6a-b, F_4,52_ = 1.03, p = 0.399) nor growth rate (Supplementary Figure S6c-d, F_4,52_ = 0.28, p = 0.888) were affected by the extract treatment.

Finally, we performed choice assays using pure glucosinolate and cardenolide compounds at the concentrations found in *E. cheiranthoides* leaves. After 24h of feeding by *P. xylostella*, leaf damage was unaffected by compound concentrations or identity (Supplementary Figure S7; leaf age: F_3,233_ = 0.97, p = 0.406; leaf age × compound: F_9,233_ = 0.148, p = 0.998).

## Discussion

The annual plant *E. cheiranthoides* produces two distinct types of chemical defences and exhibits substantial variation in these and other defence-related traits within-plant over its early development. At four weeks of age, plants were near peak vegetative growth and showed a relatively homogeneous distribution of defensive chemicals across their leaves, even though some compounds, including the most abundant cardenolide and glucosinolate, already showed a clear gradient of increasing concentrations from base to tip. Two weeks later, six-week-old plants had begun shifting to reproductive growth, indicated by the initiation of flower buds. Consequently, base-to-tip gradients strengthened in most defensive traits, both due to decreasing concentrations in oldest leaves and substantial increases in the youngest leaves.

Corresponding to the gradients in defence, we found a gradient in foliar protein concentrations of six-week-old plants, with youngest leaves containing nearly twice as much protein as the oldest leaves. In C3 plants, the photosynthesis-involved enzyme rubisco accounts for up to 65% of soluble leaf proteins (Ellis 1979), thus younger leaves on *E. cheiranthoides* not only contain a substantial fraction of total plant nitrogen, but likely also represent the most efficient photosynthetic tissues. The relative value of leaves is continuously changing throughout plant development, but the youngest leaves are generally considered to be the most valuable (Harper 1989). Younger leaves tend to be positioned highest within a plant’s canopy where they are least affected by self-shading, and young leaves usually have the highest photosynthetic rates of all leaves on a plant (Bielczynski *et al*. 2017). In consequence, the loss of younger leaves is generally more detrimental for plant fitness than loss of older leaves (Barto & Cipollini 2005).

Leaf value is further determined by the growth stage of the plant, particularly for annual plants which are limited to a single vegetation period. For these plants, theoretical models predict an abrupt and complete shift from vegetative to reproductive growth (Cohen 1971, 1976), which matches experimental findings of strong reductions in vegetative growth of annual plants after initiation of flowering (Guilbaud *et al*. 2015). Whereas a plant in its vegetative growth phase can relatively easily replace lost leaf tissue, any leaves lost on a flowering plant are thus expected to have substantially more severe fitness costs. Indeed, young leaves on *Arabidopsis thaliana* plants at the transition to flowering were found to be the most valuable for seed production (Barto & Cipollini 2005). Taken together, our results thus match the predictions of ODT (McKey 1974), with a weaker gradient in defence allocation for vegetative four-week-old plants, which then transitioned to a very steep defence gradient corresponding to the increasing value of younger leaves in early-reproductive six-week-old plants.

Gradients in defence were particularly apparent in six-week-old plants due to the significant reduction of defence concentrations in older leaves. This reduction in defence was accompanied by substantial changes in carbohydrate content, where we found the oldest leaves to have depleted starch reserves but still contain high concentrations of soluble monosaccharides. High concentrations of free sugars are a characteristic trait of senescing leaves (Wingler & Roitsch 2008), and indeed, low protein content and an observed lack of green colouration of the oldest leaves on six-week-old plants suggests that these leaves were in the process of nutrient re-mobilization and senescence. The very low concentration of defensive compounds thus was most likely the result of re-mobilization and transport as well. Glucosinolates are known to be substantially decreased in senescing leaves of *A. thaliana* (Brown *et al*. 2003), and there is active transport of these compounds via the phloem from mature leaves to inflorescences and fruits (Brudenell *et al*. 1999; Chen *et al*. 2001). Cardenolides have likewise shown to be transported from mature leaves to other tissues (Alani *et al*. 2021), with our results suggesting almost complete re-mobilization of these compounds during leaf senescence as a potential cost saving strategy.

Larvae of the diamondback moth *P. xylostella* remain one of a small set of herbivores that continue to feed on *E. cheiranthoides* under natural conditions, despite the plant’s novel cardenolide defence (Mertens *et al*. 2021; Wang *et al*. 2023). Here, we found a strong preference of caterpillars towards the oldest leaves on intact plants, particularly on six-week-old plants. Larvae of *P. xylostella* feeding on *E. cheiranthoides* are easily disturbed and frequently drop off the plant, perhaps due to its inferior quality as a host plant. Caterpillars crawling back up the plant would therefore first encounter the lower leaves and may choose to feed there for convenience rather than preference. However, at least for six-week-old plants, we could confirm preference for and increased performance on older leaves using both no-choice assays on whole plants and choice assays on detached leaves.

Given the low protein concentrations of older *E. cheiranthoides* leaves, the observed preference for and improved performance on older leaves is unlikely a result of nutritional quality. Indeed, on other Brassicaceae hosts, *P. xylostella* larvae consistently feed on the youngest, most protein-rich leaves available (Moreira *et al*. 2016). We therefore expected caterpillar preference and performance to be directly affected by the substantially increased levels of defensive traits in younger leaves. In leaves of *E. cheiranthoides*, we found concentrations of cardenolides and glucosinolates to co-vary with trichome density, thus a separation of effects in natural plants is challenging. By using a novel *E. cheiranthoides* CRISPR/Cas9 knockout mutant that lacks cardenolides (Younkin *et al*. 2024), we attempted to causally link the observed herbivore preference to cardenolide concentrations, but surprisingly, herbivore preference for older leaves was unaffected by the removal of these defences. In further support of this negative result, addition of the defences to broccoli leaf discs, either by applying crude solvent extracts from young leaves, or even pure compounds at concentrations found in young leaves, failed to repel feeding by *P. xylostella*. It thus appears that this herbivore is largely immune to variation in the cardenolide defences of *E. cheiranthoides*.

Larvae of *P. xylostella* feed on a diversity of glucosinolate-defended Brassicacae plants, and their co-evolved molecular adaptations enable these herbivores to efficiently disarm glucosinolate defences (Ratzka *et al*. 2002). High concentrations of glucosinolates in fact more commonly act as feeding stimulants for *P. xylostella* in other plant species (Badenes-Pérez *et al*. 2020). In contrast, *Erysimum* plants are the only host on which *P. xylostella* may encounter cardenolide defences, and given these plant’s sparse distribution globally, selection for specialized resistance is unlikely to be strong. Whereas many herbivores that feed on cardenolide-producing plants express cardenolide-resistant Na^+^/K^+^-ATPases in their nerve cells (target-site insensitivity, Dobler *et al*. 2012), such adaptations appear to be absent in the *P. xylostella* genome (Ward *et al*. 2021). Nonetheless, our results suggest that *P. xylostella* can cope with cardenolides when they are present in its food, perhaps due to less specific resistance mechanisms such as cardenolide diffusion barriers surrounding the insect nerve cord (Petschenka *et al*. 2013). General resistance mechanisms such as these may well be responsible for the global success of *P. xylostella* as an agricultural pest.

Given the apparent lack of involvement of chemical defences in affecting *P. xylostella* preference, the traits preventing attack of younger leaves in whole plant and cut leaf assays remain unknown. Even though *P. xylostella* has a high tolerance of glucosinolate defences due to its efficient detoxification mechanism (Ratzka *et al*. 2002), its larvae can be susceptible to glucosinolate breakdown products under some conditions, and larval growth was demonstrated to be reduced on tissues with high myrosinase activity (Li *et al*. 2000). While we tested for an effect of varying glucosinolate concentrations in our pure compound assays, we did not manipulate myrosinase activity, and such activity is known to vary with plant development (Iversen & Baggerud 1980) and may well vary with leaf age as well. Likewise, physical traits such as trichomes may also play a role in herbivore feeding preference, as these showed an equivalent vertical gradient with chemical defences. While trichomes on leaves of *Arabidopsis lyrata* were considered to be an unlikely factor in resistance against *P. xylostella*(Puentes & Ågren 2013), trichomes of *E. cheiranthoides* are substantially more dense and might therefore be more relevant. With the development of new CRISPR/Cas9 mutants, causal evaluation of glucosinolates, myrosinase activity, and trichomes in the defence of *E. cheiranthoides* may be feasible soon. In the meantime, it is clear that *E. cheiranthoides* co-expresses several measurable and perhaps additional unmeasured traits to form an elaborate defence syndrome (Agrawal & Fishbein 2006), with its highest expression in the youngest, most valuable leaves.

Cardenolides are potent chemical defences in other plant systems (Agrawal *et al*. 2012), and in *E. cheiranthoides* they are causally linked to the complete avoidance of the plant by several herbivore species (Younkin *et al*. 2024). As part of its defence syndrome, *E. cheiranthoides* co-expresses cardenolide compounds with glucosinolates and trichomes, and appears to accurately tailor expression of the novel defence to match the relative value of different organs, including an eventual re-mobilization of these defences to floral tissues and seeds (Alani *et al*. 2021). Interestingly, we discovered distinct patterns in the regulation of individual cardenolide compounds, which match the compound-specific regulation of glucosinolate compounds during plant maturation and flowering in other systems (Strauss *et al*. 2004). While we currently lack an understanding of the importance of structural variation among cardenolides of *E. cheiranthoides*, evidence from other plant systems suggests that such variation may indeed be adaptive against some herbivores (Petschenka *et al*. 2018).

In conclusion, we found that within-plant variation of a novel chemical defence mirrors that of other chemical and physical traits to prioritize defence of the most valuable plant tissues. Concerted expression and regulation of multiple costly defences within-plant thus provides additional evidence for the non-redundancy of these traits in plant defence. Even though cardenolides may not be an effective defence against all herbivore species, these compounds nonetheless appear to be important components of a carefully regulated plant defence system that provides protection against a diverse community of herbivores.

## Acknowledgments

We thank Broti Biswas and Laura Dällenbach for help with insect rearing and insect assays, and Rayko Jonas and Markus Meierhofer for help with plant production for experiments. Gordon Younkin and three anonymous reviewers provided helpful comments that improved the manuscript. We also thank Jakob Lang and Prof. Laurent Bigler of the Mass Spectrometry and General Analytical Services of the Department of Chemistry, University of Zürich, for providing us with instrument access and assistance with the analysis of plant defensive chemicals. This project has received funding from the European Research Council (ERC) under the European Union’s Horizon 2020 research and innovation programme (grant agreement No 950319). Additional support was provided by Swiss National Science Foundation grant PCEFP3_194590 to TZ.

## Data availability

All raw data and R code is accessible on Dryad (10.5061/dryad.73n5tb35b).

## Supplementary Methods

### 1. Plutella xylostella rearing

The strain of P. xylostella used in this study was originally provided by Syngenta Crop Protection AG (Münchwilen, Switzerland). We maintained this strain for continuous use in an incubator (16:8 h light: dark, 23 °C, 60% RH) on artificial diet (Beet Armyworm Diet, Frontier Scientific Services, Newark, USA) consisting of agar, wheat germ and casein, vitamins, and chlortetracycline. For continuous rearing, approximately 500 eggs of *P. xylostella* were placed in a plastic cup (250 mL volume) filled with 115 mL solidified diet. Cups were then sealed with a piece of surgical gauze and a cardboard lid. After hatching, larvae completed their entire development within the cup, before pupating on the gauze underneath the cardboard lid. The pieces of gauze with pupae attached were then removed, briefly bleached (2 min in 500 mL lukewarm water containing 10 mL 5% bleach), rinsed, and left to dry in a plastic cage (30 x 30 x 30 cm, BugDorm, MegaView Science Co., Taiwan) where adults would emerge. A 30% sugar solution on cotton wool was provided as adult food. When most adults had emerged, we fit a sheet of parafilm sprayed with cabbage juice to the inside of the cage for oviposition. Parafilm egg sheets were removed after 24h, and a subset of fresh eggs was used to set up a new generation, while the remaining eggs were immediately stored at 4 °C for up to two weeks.

### 2. Leaf extract choice and performance assays

To test the role of soluble leaf metabolites, we extracted *E. cheiranthoides* leaves and applied these extracts to broccoli leaf discs as a neutral host plant that is well-liked by *P. xylostella*. We again only used three six-week-old *E. cheiranthoides* plants to make extracts for this assay. We marked leaves of four distinct ages and collected four leaf discs (7.6 mm diameter) from each marked leaf, and extracted discs in bulk using 20 µL 70% MeOH per disc. To set up the choice assay, five broccoli leaf discs (7.6 mm diameter) were set up in a Petri dish for a total of 12 replicate dishes. In each dish, four discs were painted with 20 µL of crude extract from a different leaf age, and the remaining disc was painted with 20 µL of 70% MeOH as a control. We then added single 9-day-old *P. xylostella* larvae (Table S1) to the centre of each dish. After 24h, we extended this assay by transferring the larvae to a new, identical set of Petri dishes with painted broccoli leaf discs for an additional 24h. Leaf discs from both choice assay intervals were then scanned and quantified for area consumption.

We also performed a no-choice performance assay using single broccoli leaf discs (14.1 mm diameter) painted with extracts from one of four leaf ages of six-week-old *E. cheiranthoides* plants. We also included a fifth disc painted with 70% MeOH as a control. As with the extract choice assay, we performed extract performance assays over two 24h intervals. To generate sufficient extracts, we bulk-collected leaves from four leaf ages of six-week-old plants. Leaves were flash-frozen in liquid nitrogen, freeze-dried for 48 h, and then ground and weighed. To add an appropriate amount of crude *E. cheiranthoides* extract to broccoli leaf discs, we determined specific leaf area (SLA, mm^2^ leaf per mg dry weight) for each leaf age by collecting additional leaves on a separate set of *E. cheiranthoides* plants, determined their leaf areas by scanning, and later determined leaf dry weight (Table S3). Using SLA to determine the dry weight of one *E. cheiranthoides* leaf disc (14.1 mm diameter), we calculated the volume of extraction solvent needed to achieve crude extract solutions containing the equivalent of one leaf disc per 200 µL extraction solvent. Dried and ground leaf powder from each leaf age was extracted in bulk using 70% MeOH. Bulk extracts were then concentrated using a CentriVap vacuum concentrator (Labconco, Kansas City, USA) at 5 °C for 48h. Finally, dried extracts were resuspended by adding the equivalent of 20 µL of 70% MeOH per leaf disc (10x concentrated), followed by sonication, vortexing, and centrifugation at 2*10^4^ g for 5 min (4°C).

We filled wells of 12-well plates with a thin layer of 1% agar and added one broccoli leaf disc each well painted with 20 µL extract or 70% MeOH in a randomized pattern. We then added one pre-weighed 9-day-old larva of *P. xylostella* (Table S1) to each well (12 replicates per treatment). Plates were sealed with Parafilm to prevent the escape of caterpillars. After 24h, leaf discs were scanned for quantification of area consumed and all larvae were weighed and transferred to a new, identical set of 12-well plates with painted broccoli leaf discs. After an additional 24h, all larvae were weighed again and then discarded. Leaf discs from both assay intervals were scanned and quantified for area consumption.

**Figure S1.**
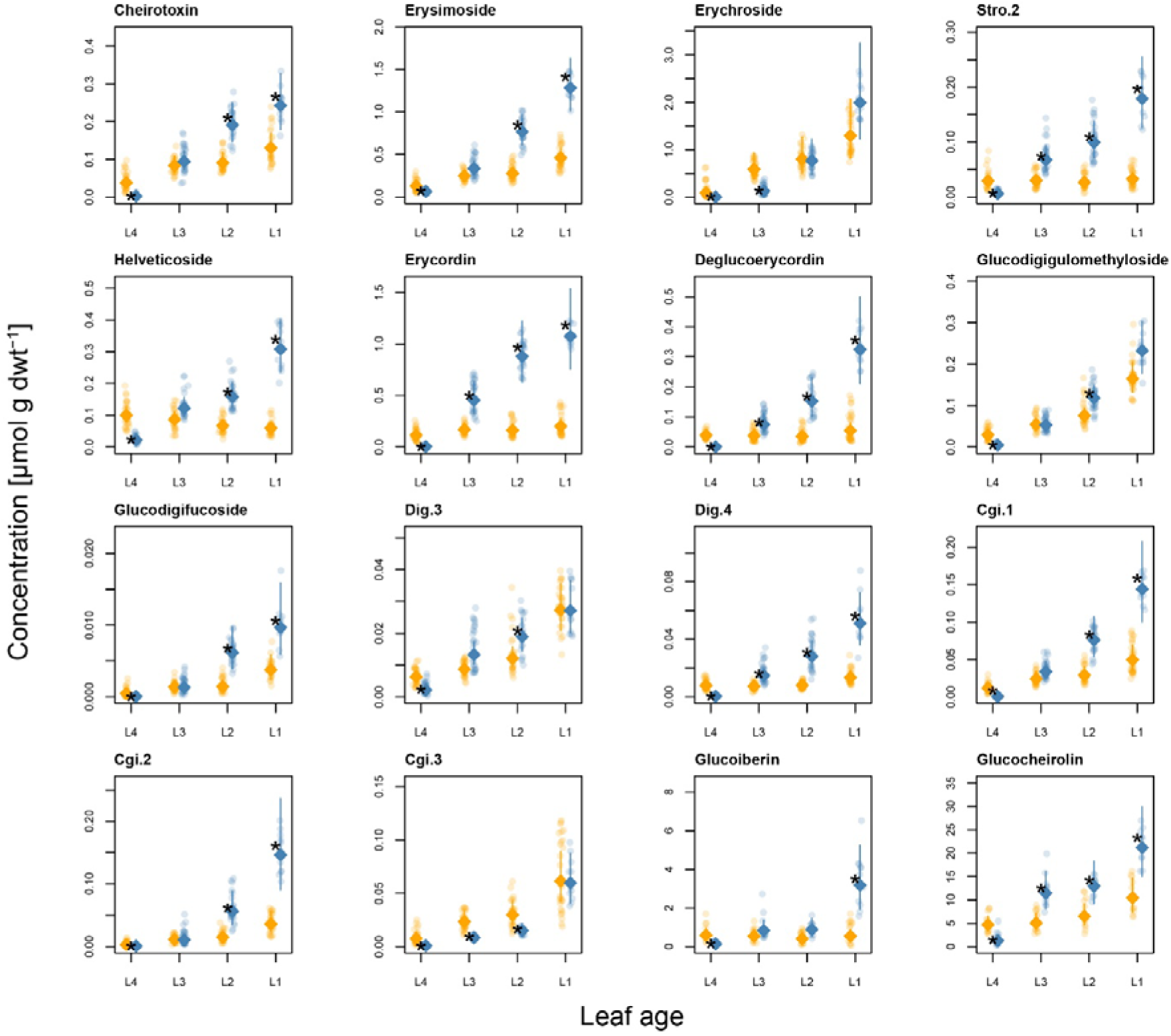
Absolute concentrations of individual defence compounds in leaves of *E. cheiranthoides* from four leaf age classes of four-week (orange symbols) and six-week-old plants (blue symbols). Points are values from individually quantified leaf discs, while diamonds and lines are the estimated mean concentrations and 95% confidence intervals from separate mixed effects models. Asterisks (*) indicate significant pairwise age differences between leaves of the same leaf age. Glucoiberin and glucocheirolin (bottom right) are the main glucosinolate compounds found in *E. cheiranthoides*, while all other compounds shown are cardenolides. Compounds Stro.2, Dig.3, Dig.4, and Cgi.1-3 are unnamed glycosides of strophanthidin, digitoxigenin, and cannogenin, respectively (see Table S2 for structural details). One identified cardenolide compound (Cgi.4, not shown) could not be quantified due to its low abundance in plant tissues.

**Figure S2.**
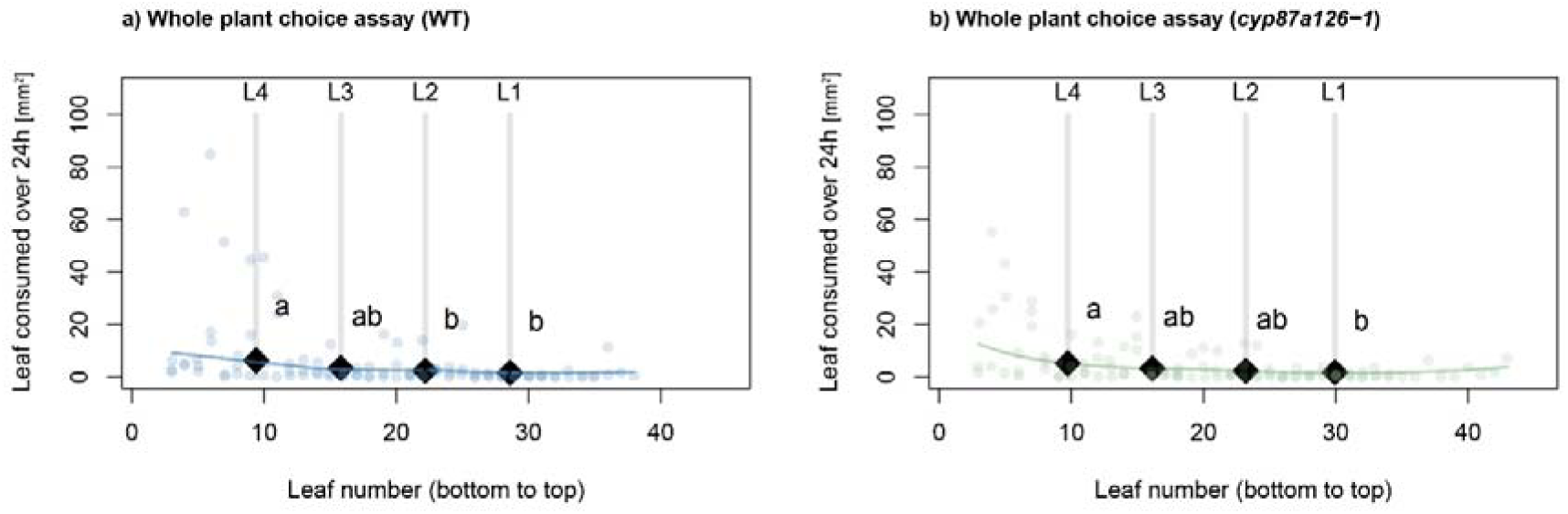
Leaf area consumption by *P. xylostella* under choice conditions on whole plants. a) Damaged leaf area (mm^2^) on six-week-old wildtype (WT) and b) cardenolide-free CRISPR/Cas9 knockout plants (n=3 each) after 24 hours of feeding by 20 free-moving caterpillars. Points are individual leaves, while coloured lines are LOESS smoothers to visualize damage patterns. Grey vertical lines and numbers indicate the position of leaf age classes used in other assays, based on leaf number percentiles (20^th^, 40^th^, 60^th^ and 80^th^ percentile). Diamonds and lines are the mean damage and 95% confidence intervals, estimated for these leaf ages from mixed effects models. Letters denote significant linear contrasts at the p<0. 05 level.

**Figure S3.**
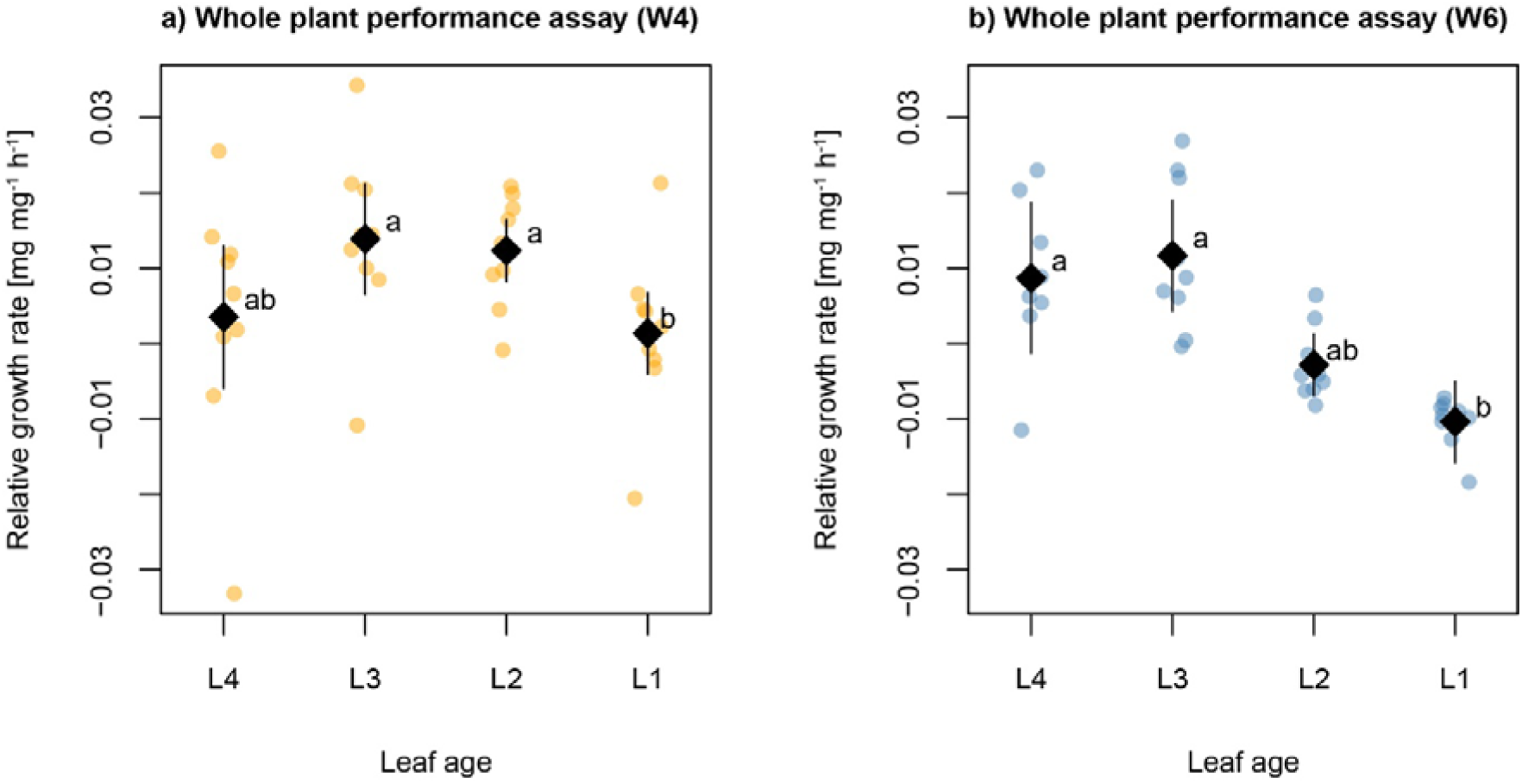
Growth rate [mg mg^-1^ h^-1^] of *P. xylostella* larvae under no-choice conditions on leaves from four age classes of a) four-week or b) six-week-old (W6) plants (n = 9 each) after 24 hours of feeding. Individual caterpillars were confined to one leaf using clip cages. Points are individual caterpillars, while diamonds and lines are the estimated mean damage areas and 95% confidence intervals from a linear model. Letters denote significant linear contrasts at the p<0.05 level.

**Figure S4.**
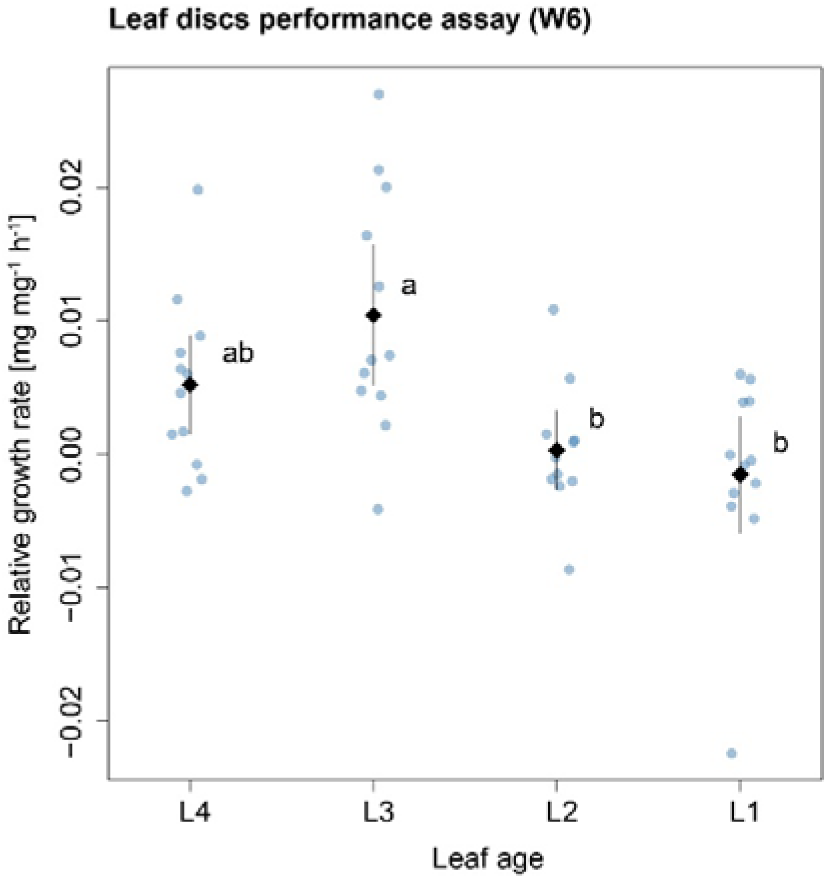
Growth rate [mg mg^-1^ h^-1^] of *P. xylostella* larvae under no-choice conditions on leaf discs from four age classes of six-week-old *E. cheiranthoides* plants (n = 12 each) after 24 hours of feeding. Points are individual caterpillars, while diamonds and lines are the estimated mean damage areas and 95% confidence intervals from a linear model. Letters denote significant linear contrasts at the p<0.05 level.

**Figure S5.**
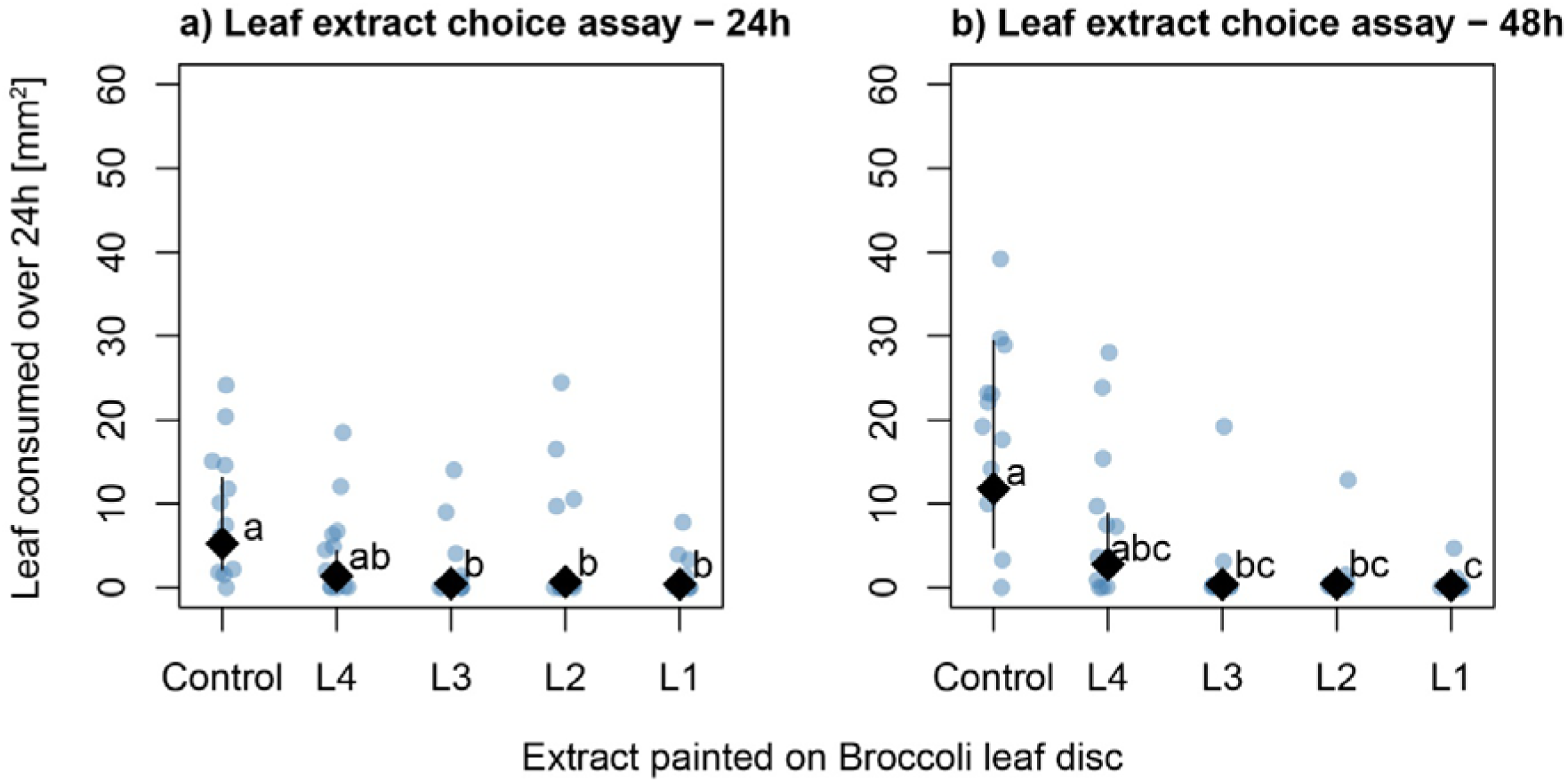
Leaf area consumption [mm^2^] by *P. xylostella* under choice conditions on detached broccoli leaf discs painted with 70% MeOH or with leaf extracts from four age classes of six-week-old *E. cheiranthoides* plants. Damaged areas on leaf discs of a five-way choice assay (n=12) using single caterpillars after a) 24 and b) 48 hours of feeding. Caterpillars were given a fresh set of leaf discs after 24 hours. Points are individual damage areas, while diamond and lines are the estimated mean damage areas and 95% confidence intervals from mixed effects models. Letters denote significant linear contrasts at the p<0.05 level.

**Figure S6.**
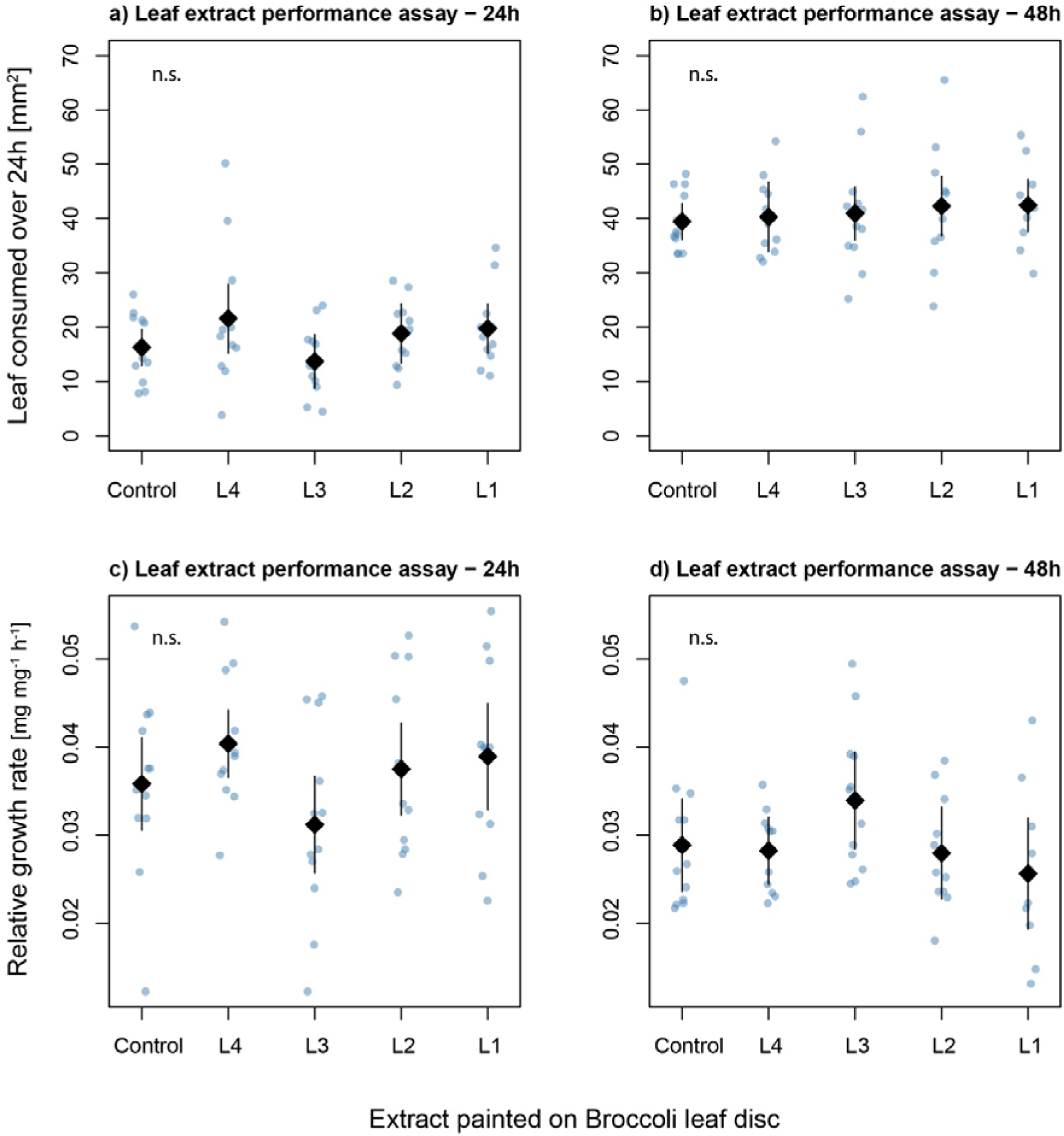
a-b) Leaf area consumption [mm^2^] and c-d) growth rate [mg mg^-1^ h^-1^] of *P. xylostella* larvae under no-choice conditions on detached broccoli leaf discs painted with 70% MeOH or with leaf extracts from four age classes of six-week-old *E. cheiranthoides* plants (n = 12 each). Caterpillars were given a fresh leaf disc after 24 hours. Points are individual leaves/caterpillars, while diamonds and lines are the estimated mean damage areas and 95% confidence intervals from a linear model. Letters denote significant linear contrasts at the p<0.05 level.

**Figure S7.**
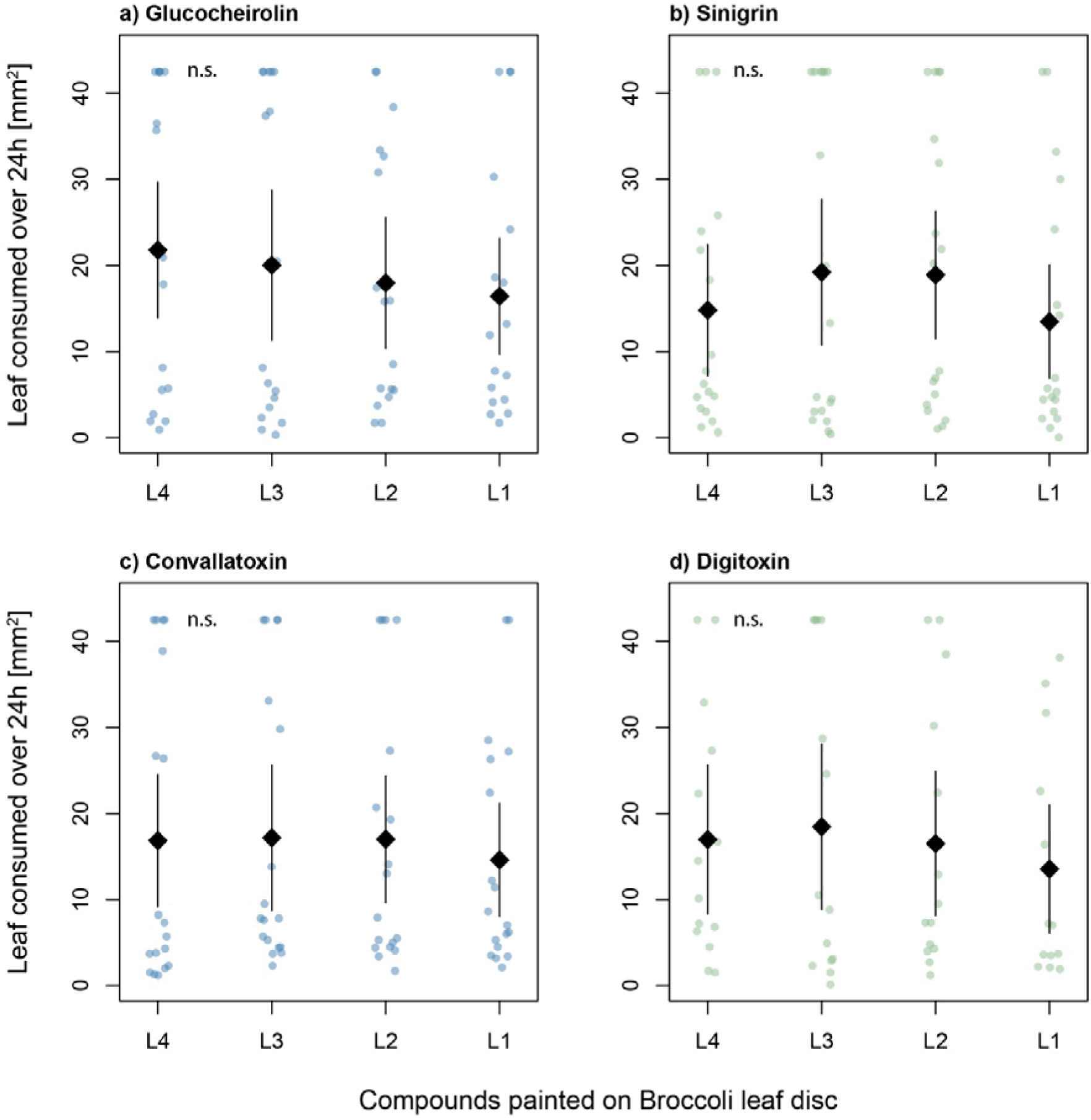
Leaf area consumption [mm^2^] by *P. xylostella* under choice conditions on detached broccoli leaf discs painted with pure compounds at concentrations corresponding to leaves of four age classes of six-week-old *E. cheiranthoides* plants. a-b) Glucosinolates glucocheirolin (occurring in *E. cheiranthoides*) and reference compound sinigrin, both added at concentrations 0.040, 0.358, 0.466, and 0.780 nmol per mm^2^ leaf area for L4-L1. c-d) Cardenolides convallatoxin (structural analogue to *E. cheiranthoides* cardenolides) and reference compound digitoxin, both added at concentrations 0.003, 0.044, 0.113, and 0.197 nmol per mm^2^ leaf area for L4-L1. Points are individual damage areas, while diamond and lines are the estimated mean damage areas and 95% confidence intervals from mixed effects models.

**Table S1.** Ages and weights of *P. xylostella* larvae used in experiments.

| Experiment | Eggs removed from rearing/fridge | Egg origin | Start of experiment | Larval age [days] | Larval stage | Larval weight range [mg] |
| --- | --- | --- | --- | --- | --- | --- |
| 1. Whole plant choice assay | 25.10.2021 | Fridge | 04.11.2021 | 10 | L3 | Not measured |
| 2. Whole plant performance assay | 25.10.2021 | Fridge | 04.11.2021 | 10 | L3 | 1.63-2.90 |
| 3. Leaf disc choice assay | 22.04.2023 | Fresh | 02.05.2023 | 10 | L3 | Not measured |
| 4. Leaf disc performance assay | 23.04.2023 | Fresh | 03.05.2023 | 10 | L3 | 1.74-2.29 |
| 5. Leaf extract choice assay | 25.09.2023 | Fridge | 03.10.2023 | 8 | L2 | 0.33-0.42 |
| 6. Leaf extract performance assays | 07.05.2024 | Fridge | 15.05.2024 | 8 | L2 | 0.33-0.42 |
| 7. Pure compound choice assay | 27.09.2024 | Fresh | 06.10.2024 | 9 | L3 | 1.71 – 2.36 |

**Table S2.**
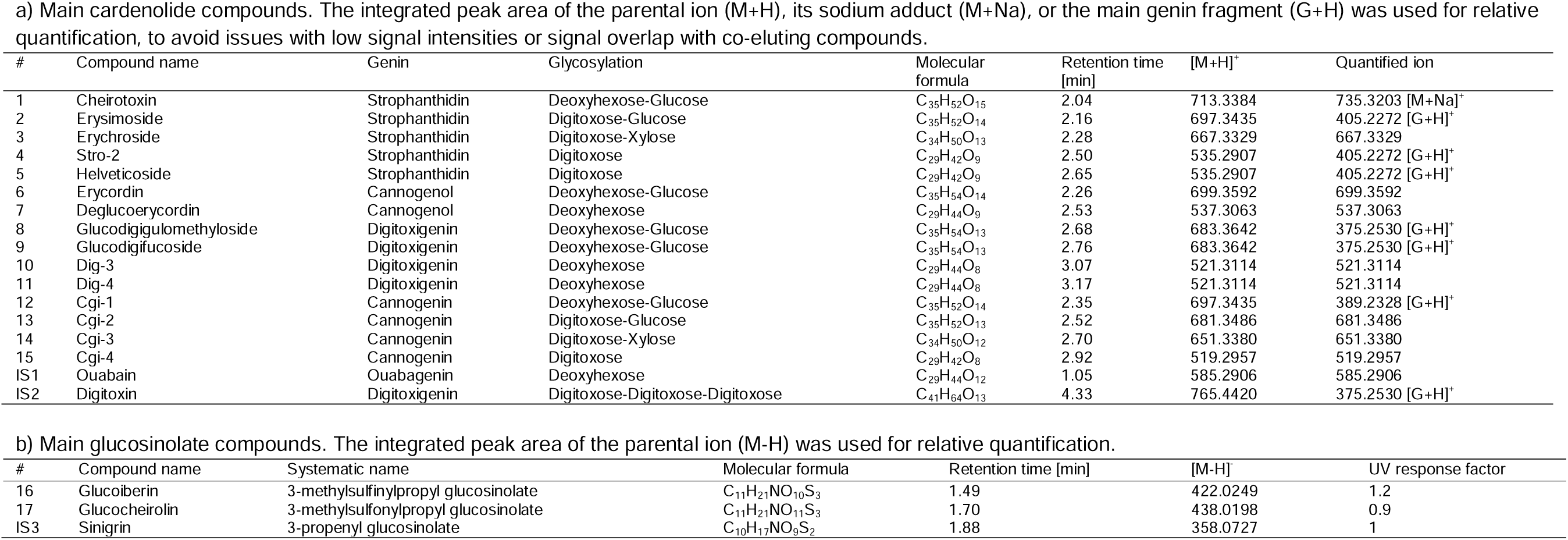
List of the main defensive metabolites produced by *E. cheiranthoides*, identified by exact mass, fragmentation patterns, and retention time. Compounds listed here are sufficiently concentrated in plant samples for reliable quantification by HPLC-UV. Other cardenolide and glucosinolate compounds were only present in trace amounts, and due to their low concentrations are less likely to be biologically relevant for plant defence.

**Table S3.** Specific leaf area (mm^2^/mg) of leaves from four leaf ages of six-week-old *E. cheiranthoides* leaves.

| Leaf age | Area (mm<sup>2</sup>) | Number of Leaves (pooled) | Weight dry (mg) | SLA (mm<sup>2</sup>/mg) |
| --- | --- | --- | --- | --- |
| L1 | 1943.8 | 5 | 62.39 | 31.16 |
| L2 | 6247.378 | 5 | 207.38 | 30.13 |
| L3 | 12507.987 | 5 | 349.27 | 35.81 |
| L4 | 10475.599 | 5 | 226.22 | 46.31 |

## References

Agrawal A., Fishbein M. (2006) Plant defense syndromes. Ecology 87:S132–S149.

Agrawal A.A., Hastings A.P. (2019) Trade-offs constrain the evolution of an inducible defense within but not between plant species. Ecology 100:e02857.

Agrawal A.A., Petschenka G., Bingham R.A., Weber M.G., Rasmann S. (2012) Toxic cardenolides: chemical ecology and coevolution of specialized plant–herbivore interactions. New Phytologist 194:28–45.

Alani M.L., Younkin G.C., Mirzaei M., Kumar P., Jander G. (2021) Acropetal and basipetal cardenolide transport in *Erysimum cheiranthoides* (wormseed wallflower). Phytochemistry 192:112965.

Badenes-Pérez F.R., Gershenzon J., Heckel D.G. (2020) Plant glucosinolate content increases susceptibility to diamondback moth (Lepidoptera: Plutellidae) regardless of its diet. Journal of Pest Science 93:491–506.

Barto E.K., Cipollini D. (2005) Testing the optimal defense theory and the growth-differentiation balance hypothesis in *Arabidopsis thaliana*. Oecologia 146:169–178.

Bekaert M., Edger P.P., Hudson C.M., Pires J.C., Conant G.C. (2012) Metabolic and evolutionary costs of herbivory defense: systems biology of glucosinolate synthesis. New Phytologist 196:596–605.

Bidart-Bouzat M.G., Kliebenstein D.J. (2008) Differential levels of insect herbivory in the field associated with genotypic variation in glucosinolates in *Arabidopsis thaliana*. Journal of Chemical Ecology 34:1026–1037.

Bielczynski L.W., Łącki M.K., Hoefnagels I., Gambin A., Croce R. (2017) Leaf and plant age affects photosynthetic performance and photoprotective capacity. Plant Physiology 175:1634–1648.

Brown P.D., Tokuhisa J.G., Reichelt M., Gershenzon J. (2003) Variation of glucosinolate accumulation among different organs and developmental stages of *Arabidopsis thaliana*. Phytochemistry 62:471–481.

Brudenell A.J.P., Griffiths H., Rossiter J.T., Baker D.A. (1999) The phloem mobility of glucosinolates. Journal of Experimental Botany 50:745–756.

Chen S., Petersen B.L., Olsen C.E., Schulz A., Halkier B.A. (2001) Long-distance phloem transport of glucosinolates in Arabidopsis. Plant Physiology 127:194–201.

Clissold F.J., Sanson G.D., Read J. (2006) The paradoxical effects of nutrient ratios and supply rates on an outbreaking insect herbivore, the Australian plague locust. Journal of Animal Ecology 75:1000–1013.

Cohen D. (1971) Maximizing final yield when growth is limited by time or by limiting resources. Journal of Theoretical Biology 33:299–307.

Cohen D. (1976) The optimal timing of reproduction. The American Naturalist 110:801–807.

Dobler S., Dalla S., Wagschal V., Agrawal A.A. (2012) Community-wide convergent evolution in insect adaptation to toxic cardenolides by substitutions in the Na,K-ATPase. Proceedings of the National Academy of Sciences 109:13040–13045.

Ellis R.J. (1979) The most abundant protein in the world. Trends in Biochemical Sciences 4:241–244.

Gong B., Zhang G. (2014) Interactions between plants and herbivores: a review of plant defense. Acta Ecologica Sinica 34:325–336.

Grosser K., van Dam N.M. (2017) A Straightforward Method for Glucosinolate Extraction and Analysis with High-pressure Liquid Chromatography (HPLC). Journal of Visualized Experiments:1–9.

Guilbaud C.S.E., Dalchau N., Purves D.W., Turnbull L.A. (2015) Is ‘peak N’ key to understanding the timing of flowering in annual plantsC? New Phytologist 205:918– 927.

Halkier B.A., Gershenzon J. (2006) Biology and biochemistry of glucosinolates. Annual review of plant biology 57:303–333.

Harper J.L. (1989) The value of a leaf. Oecologia 80:53–58.

Iversen T.-H., Baggerud C. (1980) Myrosinase activity in differentiated and undifferentiated plants of Brassicaceae. Zeitschrift für Pflanzenphysiologie 97:399–407.

Jeschke V., Kearney E.E., Schramm K., Kunert G., Shekhov A., Gershenzon J., Vassão D.G. (2017) How glucosinolates affect generalist lepidopteran larvae: growth, development and glucosinolate metabolism. Frontiers in Plant Science 8:1995.

Keith R.A., Mitchell-Olds T. (2017) Testing the optimal defense hypothesis in nature: variation for glucosinolate profiles within plants. PLoS ONE 12:e0180971.

Koricheva J. (2002) Meta-analysis of sources of variation in fitness costs of plant antiherbivore defenses. Ecology 83:176–190.

Lenth R.V. (2022) emmeans: estimated marginal means, aka least-squares means. [online] URL: https://CRAN.R-project.org/package=emmeans

Li Q., Eigenbrode S., Stringam G., Thiagarajah M. (2000) Feeding and growth of *Plutella xylostella* and *Spodoptera eridania* on *Brassica juncea* with varying glucosinolate concentrations and myrosinase activities. Journal of Chemical Ecology 26:2401– 2419.

Machado R.A.R., Ferrieri A.P., Robert C.A.M., Glauser G., Kallenbach M., Baldwin I.T., Erb M. (2013) Leaf-herbivore attack reduces carbon reserves and regrowth from the roots via jasmonate and auxin signaling. New Phytologist 200:1234–1246.

Mauricio R. (1998) Costs of resistance to natural enemies in field populations of the annual plant *Arabidopsis thaliana*. The American Naturalist 151:20–28.

Mauricio R., Rausher M.D. (1997) Experimental manipulation of putative selective agents provides evidence for the role of natural enemies in the evolution of plant defense. Evolution 51:1435–1444.

Mazerolle M.J. (2020) AICcmodavg: model selection and multimodel inference based on (Q)AIC(c). [online] URL: https://CRAN.R-project.org/package=AICcmodavg

McCall A.C., Fordyce J.A. (2010) Can optimal defence theory be used to predict the distribution of plant chemical defences? Predicting the distribution of plant chemical defences. Journal of Ecology 98:985–992.

McKey D. (1974) Adaptive patterns in alkaloid physiology. The American Naturalist 108:305– 320.

Mertens D., Bouwmeester K., Poelman E.H. (2021) Intraspecific variation in plant-associated herbivore communities is phylogenetically structured in Brassicaceae. Ecology Letters 24:2314–2327.

Moreira L.F., Teixeira N.C., Santos N.A., Valim J.O.S., Maurício R.M., Guedes R.N.C., Oliveira M.G.A., Campos W.G. (2016) Diamondback moth performance and preference for leaves of *Brassica oleracea* of different ages and strata. Journal of Applied Entomology 140:627–635.

Petschenka G., Fei C.S., Araya J.J., Schröder S., Timmermann B.N., Agrawal A.A. (2018) Relative selectivity of cardenolides for Na+/K+-ATPases from the monarch butterfly and non-resistant insects. Frontiers in Plant Science 9:1–13.

Petschenka G., Pick C., Wagschal V., Dobler S. (2013) Functional evidence for physiological mechanisms to circumvent neurotoxicity of cardenolides in an adapted and a non-adapted hawk-moth species. Proceedings of the Royal Society B: Biological Sciences 280:20123089.

Pinheiro J.C., Bates D.M., DebRoy S., Sarkar D., R Core Team (2021) nlme: Linear and Nonlinear Mixed Effects Models.

Puentes A., Ågren J. (2013) Trichome production and variation in young plant resistance to the specialist insect herbivore *Plutella xylostella* among natural populations of *Arabidopsis lyrata*. Entomologia Experimentalis et Applicata 149:166–176.

R Core Team (2021) R: a language and environment for statistical computing.

Ratzka A., Vogel H., Kliebenstein D.J., Mitchell-Olds T., Kroymann J. (2002) Disarming the mustard oil bomb. Proceedings of the National Academy of Sciences 99:11223– 11228.

Schneider C.A., Rasband W.S., Eliceiri K.W. (2012) NIH Image to ImageJ: 25 years of image analysis. Nature Methods 9:671–675.

Shroff R., Vergara F., Muck A., Svatos A., Gershenzon J. (2008) Nonuniform distribution of glucosinolates in *Arabidopsis thaliana* leaves has important consequences for plant defense. Proceedings of the National Academy of Sciences of the United States of America 105:6196–6201.

Smith A.M., Zeeman S.C. (2006) Quantification of starch in plant tissues. Nature Protocols 1:1342–1345.

Strauss S.Y., Irwin R.E., Lambrix V.M. (2004) Optimal defence theory and flower petal colour predict variation in the secondary chemistry of wild radish. Journal of Ecology 92:132–141.

Traw M.B., Feeny P. (2008) Glucosinolates and trichomes track tissue value in two sympatric mustards. Ecology 89:763–772.

Velterop J.S., Vos F. (2001) A rapid and inexpensive microplate assay for the enzymatic determination of glucose, fructose, sucrose, L-malate and citrate in tomato (*Lycopersicon esculentum*) extracts and in orange juice. Phytochemical Analysis 12:299–304.

Wang K., van Bergen E., Züst T. (2023) Escaping herbivory through chemical novelty? Field performance and herbivore attack in the cardenolide-producing crucifer plant *Erysimum cheiranthoides*.

Ward C.M., Perry K.D., Baker G., Powis K., Heckel D.G., Baxter S.W. (2021) A haploid diamondback moth (*Plutella xylostella* L.) genome assembly resolves 31 chromosomes and identifies a diamide resistance mutation. Insect Biochemistry and Molecular Biology 138:103622.

Winde I., Wittstock U. (2011) Insect herbivore counteradaptations to the plant glucosinolate-myrosinase system. Phytochemistry 72:1566–1575.

Wingler A., Roitsch T. (2008) Metabolic regulation of leaf senescence: interactions of sugar signalling with biotic and abiotic stress responses. Plant Biology 10:50–62.

Younkin G.C., Alani M.L., Páez-Capador A., Fischer H.D., Mirzaei M., Hastings A.P., Agrawal A.A., Jander G. (2024) Cardiac glycosides protect wormseed wallflower (*Erysimum cheiranthoides*) against some, but not all, glucosinolate-adapted herbivores. New Phytologist 242:2719–2733.

Züst T., Agrawal A.A. (2017) Trade-offs between plant growth and defense against insect herbivory: an emerging mechanistic synthesis. Annual Review of Plant Biology 68:513–534.

Züst T., Joseph B., Shimizu K.K., Kliebenstein D.J., Turnbull L.A. (2011) Using knockout mutants to reveal the growth costs of defensive traits. Proceedings of the Royal Society B: Biological Sciences 278:2598–2603.

Züst T., Mirzaei M., Jander G. (2018) *Erysimum cheiranthoides*, an ecological research system with potential as a genetic and genomic model for studying cardiac glycoside biosynthesis. Phytochemistry Reviews 17:1239–1251.

Züst T., Petschenka G., Hastings A.P., Agrawal A.A. (2019) Toxicity of milkweed leaves and latex: chromatographic quantification versus biological activity of cardenolides in 16 *Asclepias* species. Journal of Chemical Ecology 45:50–60.

Züst T., Rasmann S., Agrawal A.A. (2015) Growth–defense tradeoffs for two major anti-herbivore traits of the common milkweed *Asclepias syriaca*. Oikos 124:1404– 1415.

Züst T., Strickler S.R., Powell A.F., Mabry M.E., An H., Mirzaei M., York T., Holland C.K., Kumar P., Erb M., Petschenka G., Gómez J.-M., Perfectti F., Müller C., Pires J.C., Mueller L.A., Jander G. (2020) Independent evolution of ancestral and novel defenses in a genus of toxic plants (*Erysimum*, Brassicaceae). eLife 9:e51712.

